# Diversification of ligand preference in conserved Brassicaceae LORE immune receptors matches proteobacterial 3-hydroxy fatty acid profiles

**DOI:** 10.64898/2026.05.15.725355

**Authors:** Lin-Jie Shu, Lukas Schönberger, Luca Ziffermayer, Fan-Yu Yu, Gabriel Deslandes-Hérold, Christian Schmid, Michael Gigl, Milena Schäffer, Corinna Dawid, Stefanie Ranf

## Abstract

Pattern recognition receptors are often conserved across plant lineages, yet how ligand sensing diversifies during evolution remains poorly understood. The receptor kinase LORE mediates *Arabidopsis thaliana* immune responses to bacterial medium-chain 3-hydroxy fatty acids (mc-3-OH-FAs), but its distribution and functional variation across Brassicaceae remain largely unexplored. We combined phylogenetic analysis, species-wide immune profiling, ligand-binding assays, heterologous functional testing, and *A. thaliana lore-*1 complementation to define LORE diversification across Brassicaceae. LORE orthologs with conserved 3-OH-C10:0 binding are present in all major Brassicaceae lineages, including the basal lineage. Functional complementation confirmed mc-3-OH-FA-sensing capacity in orthologs spanning all four lineages. By contrast, chain-length preference profiles among mc-3-OH-FAs varied between species and orthologs, ranging from narrow 3-OH-C10:0-dominated profiles to broader profiles with comparable sensitivity to 3-OH-C10:0 and 3-OH-C12:0. 3-OH-FA profiling of a diverse collection of plant-associated bacteria revealed that high levels of mc-3-OH-FAs are prevalent in Proteobacteria, and that the range of mc-3-OH-FA chain lengths produced corresponds to the chain-length preferences of Brassicaceae LORE receptors. Our results establish LORE as a Brassicaceae-restricted pattern recognition receptor with conserved mc-3-OH-FA sensing but diversified chain-length preferences, and provide an evolutionary framework for studying how ligand preference can diversify within a biologically relevant ligand spectrum while core receptor function is maintained.

## INTRODUCTION

Plants rely on pattern recognition receptors (PRRs) at the cell surface to sense conserved microbe-associated molecular patterns (MAMPs) and activate pattern-triggered immunity (PTI). Most PRRs are receptor-like kinases (RLKs) or receptor-like proteins (RLPs). RLKs typically comprise an extracellular domain (ECD), a transmembrane domain (TMD) and an intracellular domain (ICD) comprising a kinase domain, whereas RLPs lack a major intracellular signalling domain. RLK/RLP families are highly expanded in land plants and control diverse biological processes, including development, abiotic stress responses, and immunity (Shiu & Bleecker, 2001a; Shiu *et al*., 2004; Jamieson *et al*., 2018; Dievart *et al*., 2020; Cai *et al*., 2024). While several PRRs are conserved across broader taxonomic groups, it remains poorly understood how ligand sensing and ligand selectivity diversify within a receptor family during evolution.

LORE (LIPOOLIGOSACCHARIDE-SPECIFIC REDUCED ELICITATION, alias SD1-29) belongs to the S-domain (SD)-RLK family, often also termed bulb-type (B-type) or GNA-type (*Galanthus nivalis* agglutinin; G-type) lectin RLKs, one of the major and evolutionarily dynamic RLK families in plants (Shiu & Bleecker, 2001b; Shiu & Bleecker, 2003; Shiu *et al*., 2004; Vaid *et al*., 2012; Xing *et al*., 2013; Liu *et al*., 2018; Teixeira *et al*., 2018; Dievart *et al*., 2020). In *Arabidopsis thaliana*, LORE mediates chain-length-specific immune responses to bacterial 3-hydroxy fatty acids (3-OH-FAs) and their derivatives 3-hydroxyalkanoates (HAAs): medium-chain (mc)-3-OH-FAs (C8 to C12) activate immunity, whereas longer-chain 3-OH-FAs (C14 and C16) are inactive (Ranf *et al*., 2015; Kutschera *et al*., 2019; Schellenberger *et al*., 2021). Within the active medium-chain range, ligand sensitivity also differs with chain length: 3-hydroxydecanoic acid (3-OH-C10:0) activates the strongest response, and activity decreases at both longer and shorter chain lengths in a stepwise manner (Kutschera *et al*., 2019). In addition, free 3-OH-FAs are the strongest elicitors, and immunogenic activity depends on structural features such as hydroxyl position and configuration, as well as carboxyl group modification (Kutschera *et al*., 2019). Mechanistically, LORE requires homomerisation via its ECD and TMD as well as kinase activity for receptor function (Luo *et al*., 2020; Wang *et al*., 2023; Eschrig *et al*., 2024b). Notably, *Arabidopsis lyrata* and *Arabidopsis halleri* encode LORE orthologs but lack detectable mc-3-OH-FA responses; their orthologs retain 3-OH-C10:0 binding yet fail to homomerize (Shu *et al*., 2021; Eschrig *et al*., 2024b). Heterologous expression experiments further showed that LORE can confer mc-3-OH-FA-responsiveness in otherwise insensitive plant species (Shu *et al*., 2021; Eschrig *et al*., 2024a; Eschrig *et al*., 2024b; Li *et al*., 2025b). Together with the identification of the Arabidopsis SD-RLK RESISTANT TO DFPM-INHIBITION OF ABSCISIC ACID SIGNALING 2 (RDA2/SD1-12) as a receptor for sphingolipid-derived molecules such as 9-methyl sphingoid base (Kato *et al*., 2022), these findings support a broader relevance of SD-RLKs in plant immunity. These features make LORE a suitable model for analysing how conserved ligand sensing can diversify in ligand selectivity and receptor functionality.

Current evidence indicates that LORE is restricted to Brassicaceae (Ranf *et al*., 2015), a family that includes the model plant *A. thaliana* as well as many crop species and for which extensive genomic and phylogenetic resources are available (The 1001 Genomes Consortium, 2016; Borsch *et al*., 2020; Winkelmüller *et al*., 2021). This provides a suitable framework to analyse receptor evolution across related species while retaining sufficient phylogenetic depth for comparative analyses. Despite the importance of LORE in *A. thaliana*, it remains unclear to what extent LORE-mediated mc-3-OH-FA sensing and chain-length selectivity are conserved across Brassicaceae, and how this relates to the bacterial mc-3-OH-FA spectrum to which plants are exposed.

Here, we used LORE as a model to examine how PRR function evolves within a plant family while ligand chemistry remains comparable. To address this, we combined comparative immune profiling across Brassicaceae with phylogenetic analysis of LORE orthologs, ligand-binding assays and functional receptor testing in heterologous and complementation systems, and related these data to 3-OH-FA profiles of plant-associated bacteria. This integrative approach allowed us to distinguish between conserved and diversified features of LORE-mediated mc-3-OH-FA sensing and to place receptor variation in a biologically relevant bacterial context.

## RESULTS

### Responses to medium-chain 3-OH-FAs vary across Brassicaceae

To explore the diversity of mc-3-OH-FA sensing across Brassicaceae, we screened leaves from 41 species for ROS responses to 3-OH-FAs of different chain lengths. The species panel covers the four major Brassicaceae lineages: basal lineage, lineage I, II (including extended lineage II), and III, as well as one species from Cleomaceae, the sister family to Brassicaceae (**Table S1**) (Huang *et al*., 2016; Kiefer *et al*., 2019; Nikolov *et al*., 2019). *A. halleri* and *A. lyrata* were analysed previously (Shu *et al*., 2021; Eschrig *et al*., 2024b), and the results are included for completeness. The non-Brassicaceae species tested, including the outgroup species *Cleome violacea*, did not respond to 3-OH-FAs (**Fig. 1A; S1, S2**). Within Brassicaceae, approximately 30% of species did not respond to any mc-3-OH-FA, but produced ROS upon flg22 elicitation, confirming functional ROS production capacity (**Fig. 1; S2**). Mc-3-OH-FA-responsive and unresponsive species co-occurred in lineages I, II, and III (**Fig. 1; S2**). Phylogenetic parsimony favours a single gain of mc-3-OH-FA-sensing capacity in the ancestor of these lineages, followed by independent losses in individual species, over repeated independent acquisitions (Cunningham *et al*., 1998). Because the only tested species from the basal lineage, *Aethionema arabicum*, did not respond to mc-3-OH-FAs, the ancestral state at the base of Brassicaceae could not be resolved at this stage (**Fig. 1A; S2**). Mc-3-OH-FA responsiveness did not show a clear association with lineage, tribe, or genus classification (Al-Shehbaz, 2012; Huang *et al*., 2016; Kiefer *et al*., 2019; Nikolov *et al*., 2019), consistent with multiple independent losses. None of the Brassicaceae species tested responded to 3-OH-C14:0 (**Fig. 1; S2**). Among 68 *A. thaliana* accessions, preselected from the 1001 Genome database to cover most available non-synonymous single-nucleotide polymorphisms (SNPs) in the At*LORE* coding sequence (CDS) (**Table S2**), only accession T740 with an L29F polymorphism in the ECD did not respond to mc-3-OH-FAs; the reason for this is under investigation (**Fig. S3**). In Col-0, LORE senses mc-3-OH-FAs in a chain length-dependent manner: 3-OH-C10:0 is the strongest agonist, and activity decreases at both shorter and longer chain lengths, with 3-OH-C13:0 and 3-OH-C14:0 eliciting no detectable response (**Fig. 1B**) (Kutschera *et al*., 2019). While the preference for 3-OH-C10:0 over 3-OH-C12:0 was largely conserved across *A. thaliana* accessions (**Fig. S3**), the mc-3-OH-FA sensitivity profiles differed markedly among Brassicaceae species. All mc-3-OH-FA-sensitive species responded strongly to 3-OH-C10:0, yet many species also showed strong responses to shorter and longer-chain mc-3-OH-FAs, including 3-OH-C8:0 and 3-OH-C13:0 (**Fig. 1B; S2**). These results demonstrate that specificity for mc-3-OH-FAs is generally conserved across Brassicaceae, while the chain-length preference pattern varies among species.

**Figure 1.**
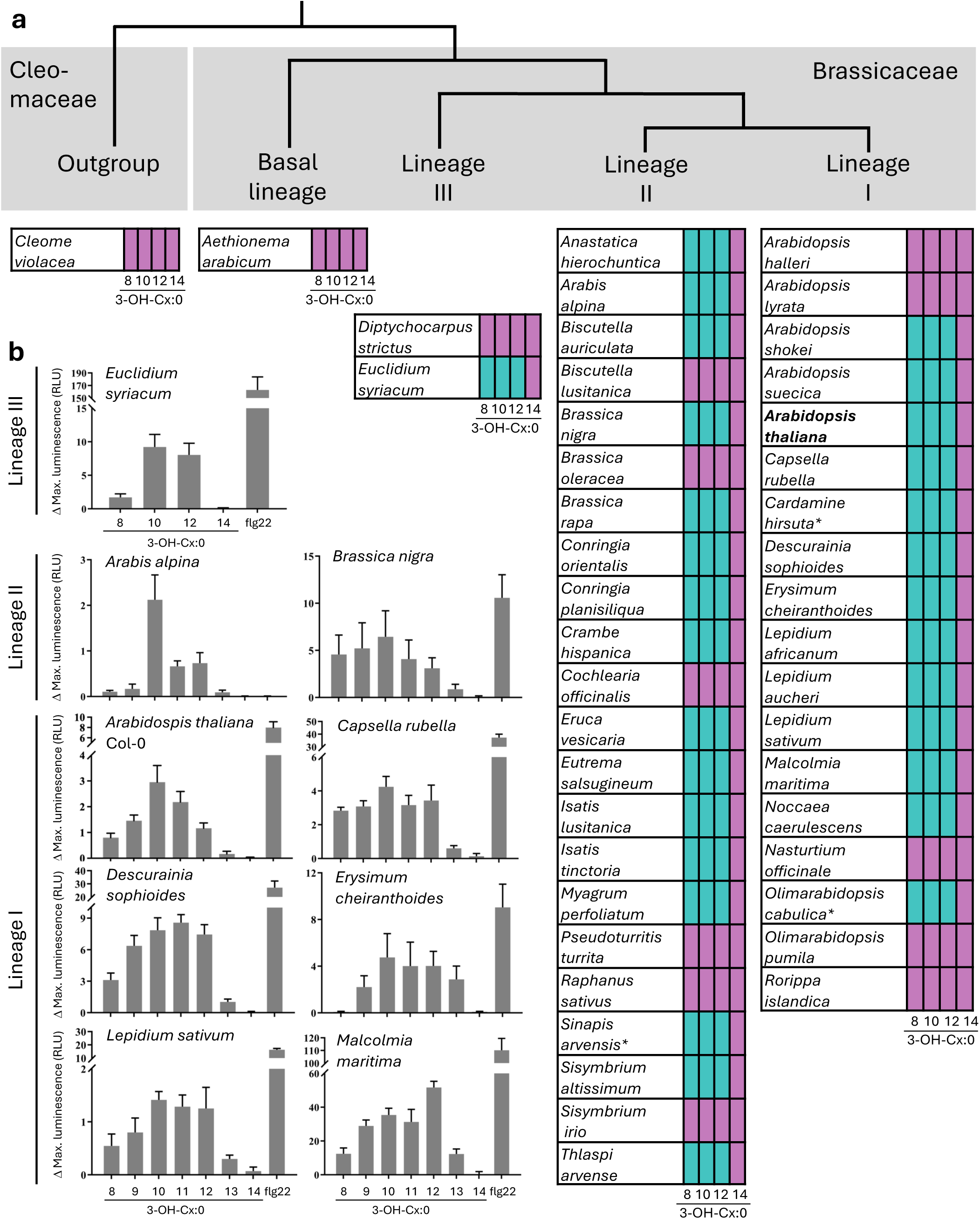
Diversity of 3-OH-FA responsiveness in Brassicaceae. **a** The tables summarise 3-OH-FA responsiveness in leaf disks of 43 species, sorted by lineage according to (Huang et al., 2016; Kiefer et al., 2019; Nikolov et al., 2019), based on 3-OH-FA-elicited ROS production shown in **b** and **Figure S2**. ROS responses of *Arabidopsis lyrata* and *A. halleri* are from Eschrig et al., 2024b. Turquoise indicates ROS response, and purple indicates no ROS response above control levels. * indicates a very low ROS response to 3-OH-FAs. **b** Examples of ROS responses to 3-OH-FAs in different species from different lineages (extended version see **Figure S2**). Maximal ROS accumulation above MeOH control level in leaf discs of the indicated Brassicaceae species upon elicitation with 5 µM 3-OH-FAs of the indicated chain length or 500 nM flg22 (mean ± SD, n = 8). Two or three independent experiments yielded similar results.

### LORE orthologs are conserved across Brassicaceae and retain 3-OH-C10:0 binding

In *A. thaliana*, *At*LORE senses mc-3-OH-FAs, whereas related SD-RLKs, such as the closest paralog, *At*SD1-23, do not (Ranf *et al*., 2015; Kutschera *et al*., 2019; Shu *et al*., 2021). Both LORE-like and SD1–23-like genes appear to have evolved specifically within Brassicaceae species (**Fig. S4**). In Brassicaceae, LORE-like receptors are presumably the primary factor facilitating mc-3-OH-FA sensing. The presence, expression patterns and functionality of these LORE-like receptors may determine the responsiveness of Brassicaceae species to mc-3-OH-FAs. We systematically investigated LORE-like receptor genes across the Brassicaceae, utilising 23 whole-genome-sequenced species spanning all four lineages (**Table S4**), including mc-3-OH-FA-responsive and unresponsive species (**Fig. 1; S2**). BLASTp identified at least one SD-RLK with ≥60% sequence identity to *At*LORE in each species. The phylogeny of the candidate sequences, supplemented with four closely related *A. thaliana* SD-RLKs, i.e., *At*SD1-23, *At*SD1-27, *At*SD1-28, and *At*SD1-30 (Vaid *et al*., 2012; Teixeira *et al*., 2018), and the more distantly related *At*RDA2/SD1-12 (Kato *et al*., 2022) as references for different outgroups (**Table S4**), was analysed. Analysis of the full-length proteins or the ligand-binding ECD yielded similar results (**Fig. 2A; S5**). The previously verified LORE ortholog from *C. rubella*, *Crub*LORE (Shu *et al*., 2021; Eschrig *et al*., 2024b), clustered with *At*LORE and other LORE-like proteins from lineage I species (**Fig. 2A; S5**). LORE orthologs from lineage I and II species clustered into separate clades, while the phylogeny of the basal lineage and lineage III was not fully resolved (**Fig. 2A; S5**). Several species, mostly from lineage II, have multiple *LORE*-like genes. While some seem to have arisen recently, based on their close phylogeny, possibly through gene duplication (Xing *et al*., 2013), others belong to separate subclades, suggesting more ancient duplication events, possibly also polyploidization (Walden *et al*., 2020) (**Fig. 2A; S5**). *At*SD1-23 was classified into a separate clade from *At*LORE (**Fig. 2A; S4, S5**). Whereas LORE-like protein sequences were found in all Brassicaceae lineages and species analysed, neither LORE-like nor SD1-23-like receptors were identified in the outgroup species *C. violacea* (**Fig. 2A; S4**), corroborating that *LORE* first appeared in Brassicaceae. Based on genome sequences, we cloned the CDSs of *LORE-*like genes from 20 species spanning all Brassicaceae lineages, and several candidates from the outgroup Cleomaceae, SD1-23-like receptors (**Fig. 3**), and *At*RDA2/SD1-12 as controls for functional analyses (**Table S2; Fig. 2A; S5**). We expressed the ECDs as apoplastic mCherry fusion proteins in *N. benthamiana* (**Fig. S6**) and tested them for binding to 3-OH-C10:0 using a ligand-depletion assay (Shu *et al*., 2021). All ECDs classified as LORE-like receptors from all four Brassicaceae lineages bound 3-OH-C10:0, including those from mc-3-OH-FA-unresponsive species (**Fig. 1; S2**), but none of the controls (**Fig. 2B**). Several species, mainly in lineage II as well as lineage III species *E. syriacum*, have multiple 3-OH-C10:0-binding LORE-orthologous ECDs (**Fig. 2**). The only LORE-like receptor ECD from the mc-3-OH-FA-unresponsive species *A. arabicum* from the basal lineage, *Aara*LORE, also bound 3-OH-C10:0 (**Fig. 2**). In conclusion, 3-OH-C10:0 binding is specific to LORE and highly conserved in Brassicaceae.

**Figure 2.**
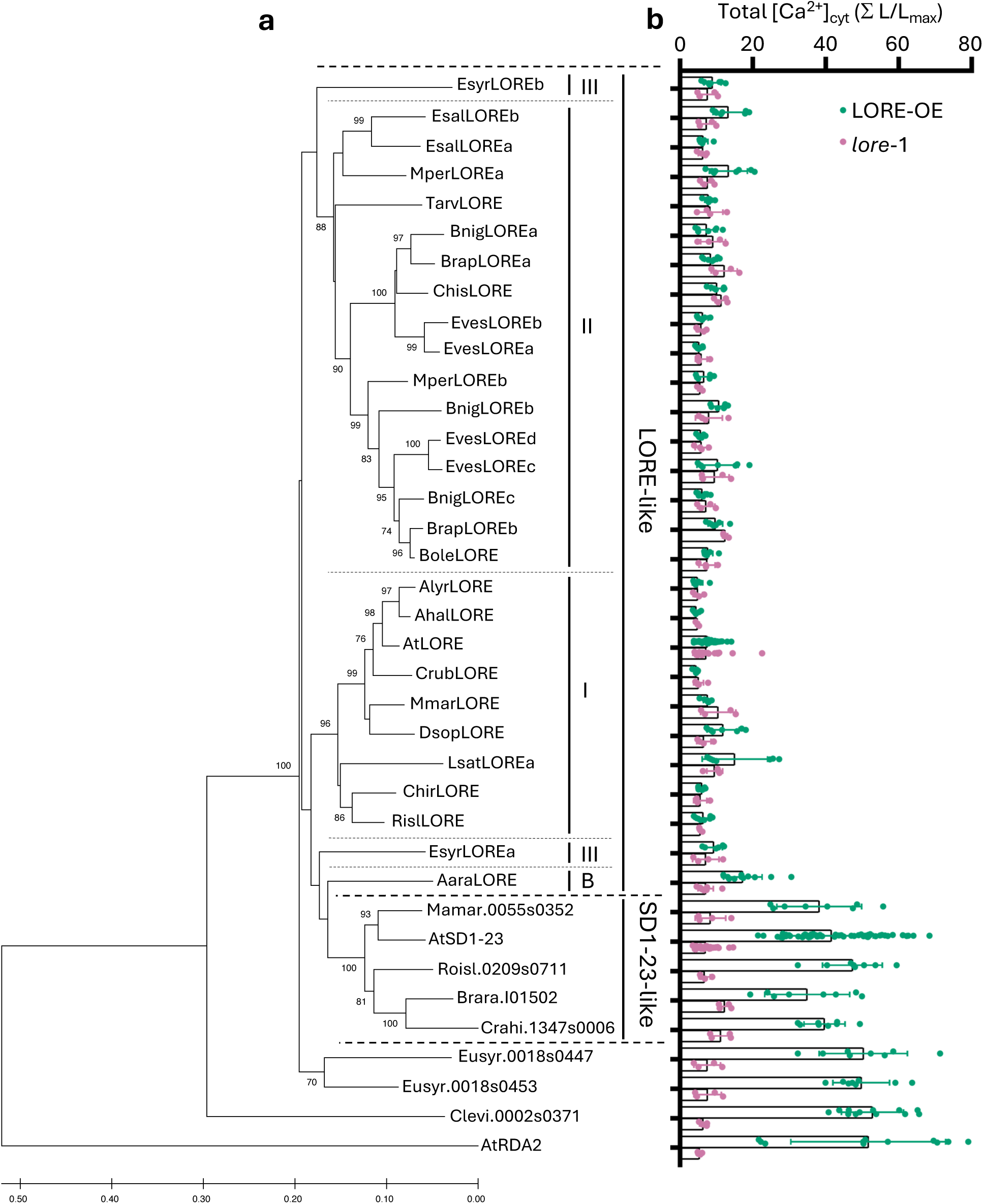
Interaction of ECDs of LORE-like receptors and other SD-RLKs with 3-OH-C10:0. **a** Phylogenetic analysis of ECDs of LORE-like receptors and other SD-RLKs. 37 protein sequences were aligned by MUSCLE. Phylogeny was analysed in MEGA XI using the Neighbour-Joining method with bootstrap support (1000 replicates). Bootstrap values over 70 are shown next to branches. Scale bar indicates evolutionary distance computed using the Poisson correction method. I, II, III, and B (basal) indicate the major lineages of the original species of the LORE-like receptors. **b** Result of ligand-depletion assay of LORE-like receptors and other SD-RLKs performed with 500 nM 3-OH-C10:0 and concentrated apoplastic washing fluids (AWFs) from *N. benthamiana* containing the indicated ECD-mCherry fusion proteins (1.5 mg/mL total protein concentration; anti-RFP immunoblots shown in **Figure S6**). Increases in [Ca^2+^]_cyt_ were measured in LORE overexpressing (LORE-OE/*lore*-1) and knockout (*lore-*1) Arabidopsis seedlings treated with filtrates from the depletion assay (mean ± SD, pooled data from at least two independent experiments for each ECD; total number of seedlings used for AtLORE-mCherry and eAtSD1-23-mCherry: LORE-OE: n=67, *lore*-1: n=34, AtSD1-12, AaraLORE-mCherry, CrubLORE-mCherry, and Clevi.0002s0371-mCherry: LORE-OE: n=12, *lore-*1: n=6; other ECDs: LORE-OE: n=8, *lore*-1: n=4).

**Figure 3.**
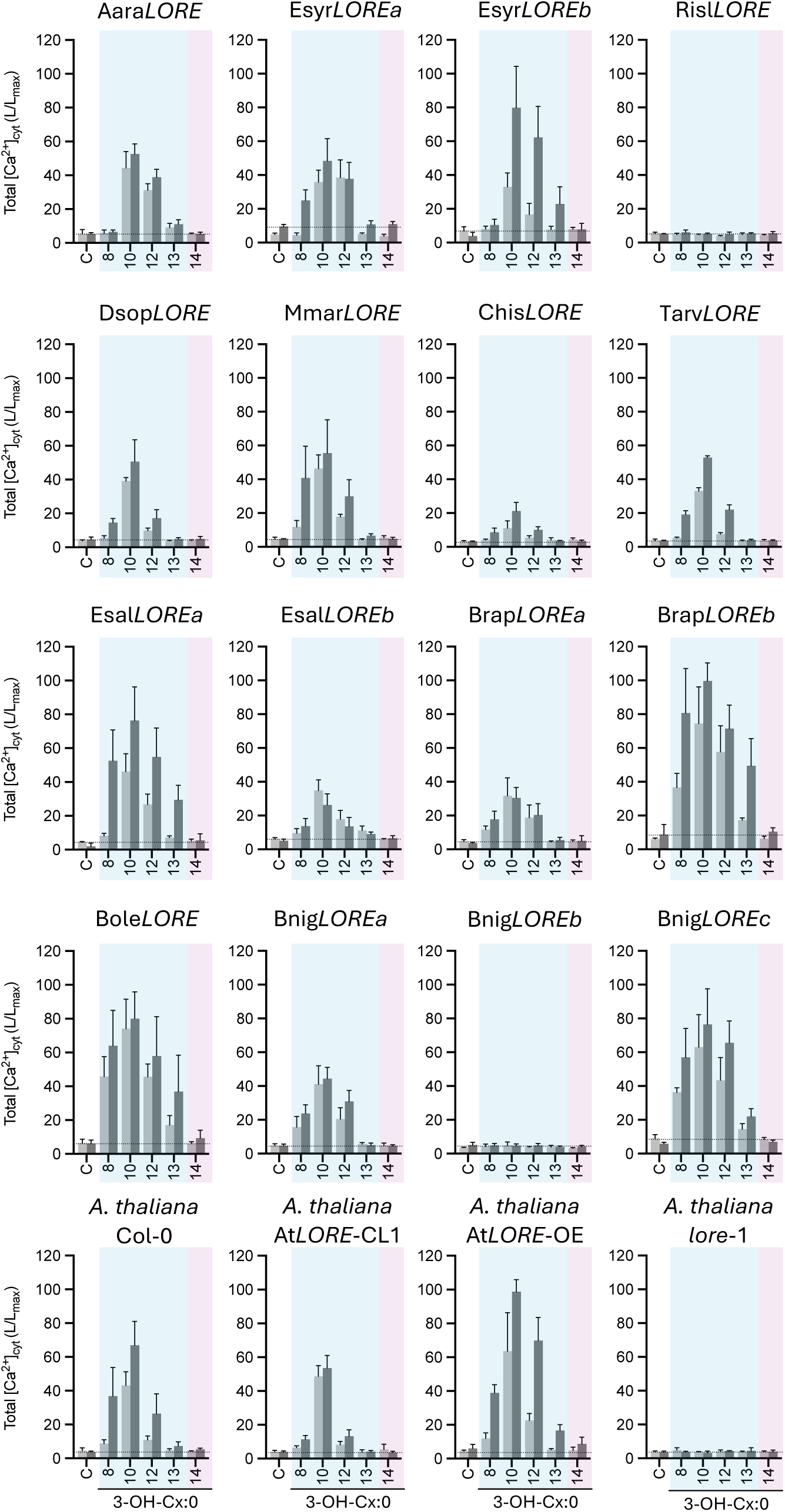
3-OH-FA chain-length preference of LORE-like receptors expressed in *Arabidopsis thaliana lore-*1. Total [Ca^2+^]_cyt_ responses were measured in *LORE* ortholog-complemented *A. thaliana lore*-1 seedlings after elicitation with 1 µM (light grey) or 5 µM (dark grey) 3-OH-FAs of the indicated chain length or MeOH as control (C). Dashed lines indicate average total [Ca^2+^]_cyt_ levels of controls as reference; shaded areas indicate medium (blue) or long (magenta) chain-length range of 3-OH-FAs. The T3 generations analysed were homozygous (based on FastRed), or, for heterozygous lines, FastRed-positive seeds were selected. One of two independent experiments with the same pattern is shown for 5 µM and one experiment for 1 µM (mean ± SD; n ≥ 4 seedlings per ortholog and chain length). One representative transgenic line from two to three independent lines analysed per ortholog is shown (see **Figure S9**). CL, complementation line; OE, overexpressing line.

### LORE orthologs differ in chain-length preference among mc-3-OH-FAs

To assess the functional capacity of individual LORE orthologs, we tested 16 orthologs from all four Brassicaceae lineages for complementation of the *A. thaliana lore-*1 knockout mutant using a previously established At*LORE* promoter fragment (Ranf *et al*., 2015; Kutschera *et al*., 2019). For each ortholog, we obtained several independent transgenic lines with validated *LORE* expression (**Fig. S7**). In most lines, mc-3-OH-FA-induced [Ca^2+^]_cyt_ signalling was restored, albeit to varying degrees (**Fig. 3; S8**). Among the orthologs tested from mc-3-OH-FA-responsive species, at least one per species restored the response to 3-OH-C10:0 in *lore-*1 (**Fig. 3; S8**). The two very closely related *Esal*LORE receptors elicited responses of different intensities to 3-OH-C10:0, while the two more distantly related *Esyr*LORE receptors elicited responses of similar intensity (**Fig. 2, 3; S5, S8**). Among orthologs from mc-3-OH-FA-unresponsive species, *Aara*LORE (basal lineage) and *Bole*LORE (lineage II) complemented *lore*-1, demonstrating that functional mc-3-OH-FA-sensing capacity is present in all major Brassicaceae clades, including the basal lineage. The phylogenetically most distant ortholog, *Aara*LORE, restored 3-OH-C10:0 signalling to levels comparable to the *A. thaliana* wild type (**Fig. 3; S5, S8**), suggesting that the core signalling mechanism has been conserved since the early diversification of Brassicaceae. By contrast, *Risl*LORE from *R. islandica* did not complement despite confirmed 3-OH-C10:0 binding capacity and transgene expression (**Fig. 2; S6, S7**), indicating that this ortholog fails to signal in the *A. thaliana* context. To dissect the chain-length sensing profiles of individual LORE orthologs, we analysed [Ca²⁺]_cyt_ responses to 3-OH-FAs of various chain lengths in the complementation lines (**Fig. 3; S9**) at two elicitor concentrations: a high concentration (5 µM) to assess the general sensing capacity under elicitor-saturating conditions, and a lower concentration (1 µM) to discern the sensing preference among different chain lengths. Even at high concentration, responses to 3-OH-C14:0 were absent or very low across all LORE orthologs, whereas several orthologs showed significant responses to 3-OH-C13:0 (e.g. *Bole*LORE and *Brap*LOREb). At the lower concentration, responses to 3-OH-C13:0 were strongly diminished. All responsive orthologs showed the strongest response to 3-OH-C10:0 at both concentrations, but the relative response to 3-OH-C8:0 and 3-OH-C12:0 varied considerably among orthologs. Because the results in the complementation lines depend on receptor levels (compare *A. thaliana* wild-type and At*LORE*-OE responses in Fig. 3) and on compatibility with the *A. thaliana* signalling network, we used a chimeric receptor approach to isolate the contribution of the ECD to chain-length preference. Chimeric receptors comprising the ECDs of different orthologs fused to the *At*LORE transmembrane and intracellular domains (ECD^Ortho^-TMD^At^-ICD^At^) were transiently expressed in *N. benthamiana* and tested at the lower elicitor concentration to resolve differences in chain-length preference (**Fig. 4**). This approach revealed a clear preference across all responsive orthologs for 3-OH-C10:0 and 3-OH-C12:0 over 3-OH-C8:0, with minimal responses to 3-OH-C13:0. Notably, orthologs differed in the relative preference for 3-OH-C10:0 versus 3-OH-C12:0: some displayed a narrow preference pattern with dominant 3-OH-C10:0 sensing, similar to *A. thaliana* Col-0, while others responded similarly to both chain lengths (**Fig. 4**). *Aara*LORE shows a distinctive profile: it similarly senses 3-OH-C10:0 and 3-OH-C12:0 but not 3-OH-C8:0, a pattern that became more pronounced at higher ligand concentrations, suggesting that LORE orthologs in other Brassicaceae species diversified from this ancestor to variants with varying degrees of specificity (**Fig. 3, 4; S9**). Overall, these results demonstrate that mc-3-OH-FA receptors have been conserved since the emergence of the Brassicaceae, but their chain-length preference profiles have diversified, with variation encoded largely in the ECD.

**Figure 4.**
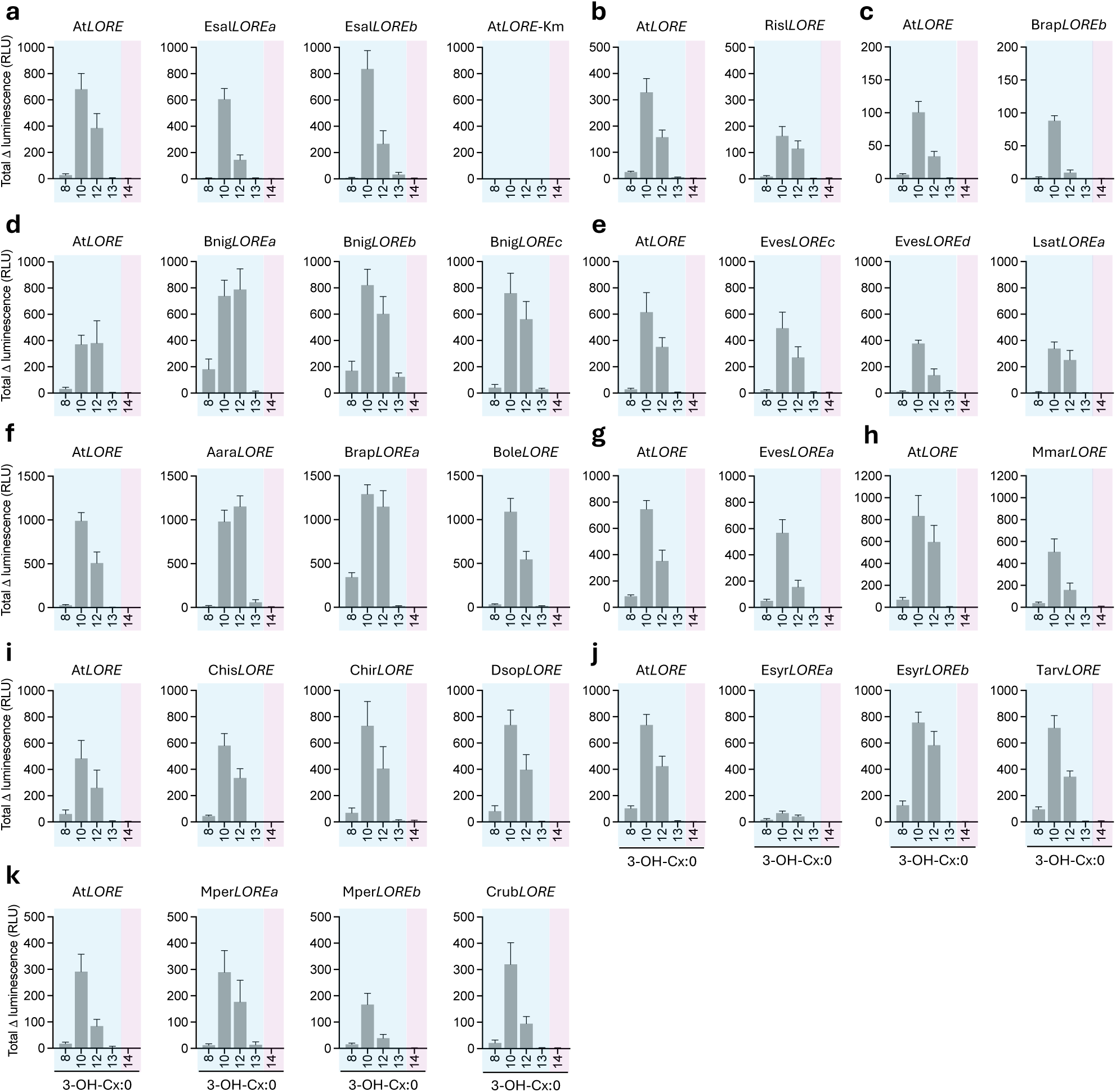
3-OH-FA chain-length preference of LORE-like receptor chimera expressed in *Nicotiana benthamiana*. **a-k** ROS production was measured 3 dpi in leaf disks from *N. benthamiana* leaves expressing *LORE* ortholog-chimera or *At*LORE as reference after elicitation with 1 µM 3-OH-FAs of the indicated chain length or MeOH as control. Three *LORE* ortholog-chimera and *At*LORE were expressed on the same leaves. Kinase-mutated *At*LORE-Km was used as a negative control. Data show mean ± SD of n = 8 leaf disks per ortholog and chain length obtained from a total of 4 replicate leaves from 2 plants. Shaded areas indicate medium (blue) and long (magenta) chain-length range of 3-OH-FAs.

### Proteobacterial 3-OH-FA profiles match the chain-length preferences of Brassicaceae LORE receptors

LORE contributes to antibacterial immunity in Arabidopsis, and LORE-mediated mc-3-OH-FA sensing was first identified using Pseudomonas and Xanthomonas (Ranf *et al*., 2015; Kutschera *et al*., 2019; Wang *et al*., 2023). To explore the biological relevance of the observed differences in 3-OH-FA chain-length preference among Brassicaceae, we profiled free 3-OH-FA content across a broad panel of plant-associated bacteria. The panel comprised a core collection of bacterial isolates from the natural *A. thaliana* leaf microbiota (Bai *et al*., 2015; Carlström *et al*., 2019) spanning Proteobacteria, Actinobacteria, Bacteroidetes, and Firmicutes, as well as pathogenic and non-pathogenic Pseudomonas isolates (Karasov *et al*., 2018) and prominent Xanthomonas pathogens (**Fig. 5; S10**). Most isolates produced only low levels of 3-OH-FAs. By contrast, high levels of elicitor-active mc-3-OH-FAs were predominantly detected in Proteobacteria, particularly in β- and γ-Proteobacteria (**Fig. 5; S10**). 3-OH-C10:0 was highly abundant in various strains, including several *P. syringae* and *P. fluorescens* isolates, while 3-OH-C12:0 predominated in *Xanthomonas* and a related *Stenotrophomonas* isolate. One *Xylophilus* isolate produced notably high amounts of 3-OH-C8:0. Importantly, none of the strains analysed showed comparably high levels of 3-OH-C14:0 (**Fig. 5; S10**). Thus, the mc-3-OH-FA chain-length range preferentially sensed by Brassicaceae LORE orthologs corresponds to the most abundant mc-3-OH-FAs produced by Proteobacteria, a major group of plant-associated Gram-negative bacteria. Given that these Proteobacteria also release relevant amounts of mc-3-OH-FAs during interactions with the plant, this provides a plausible source of LORE ligands during plant–bacteria interactions.

**Figure 5.**
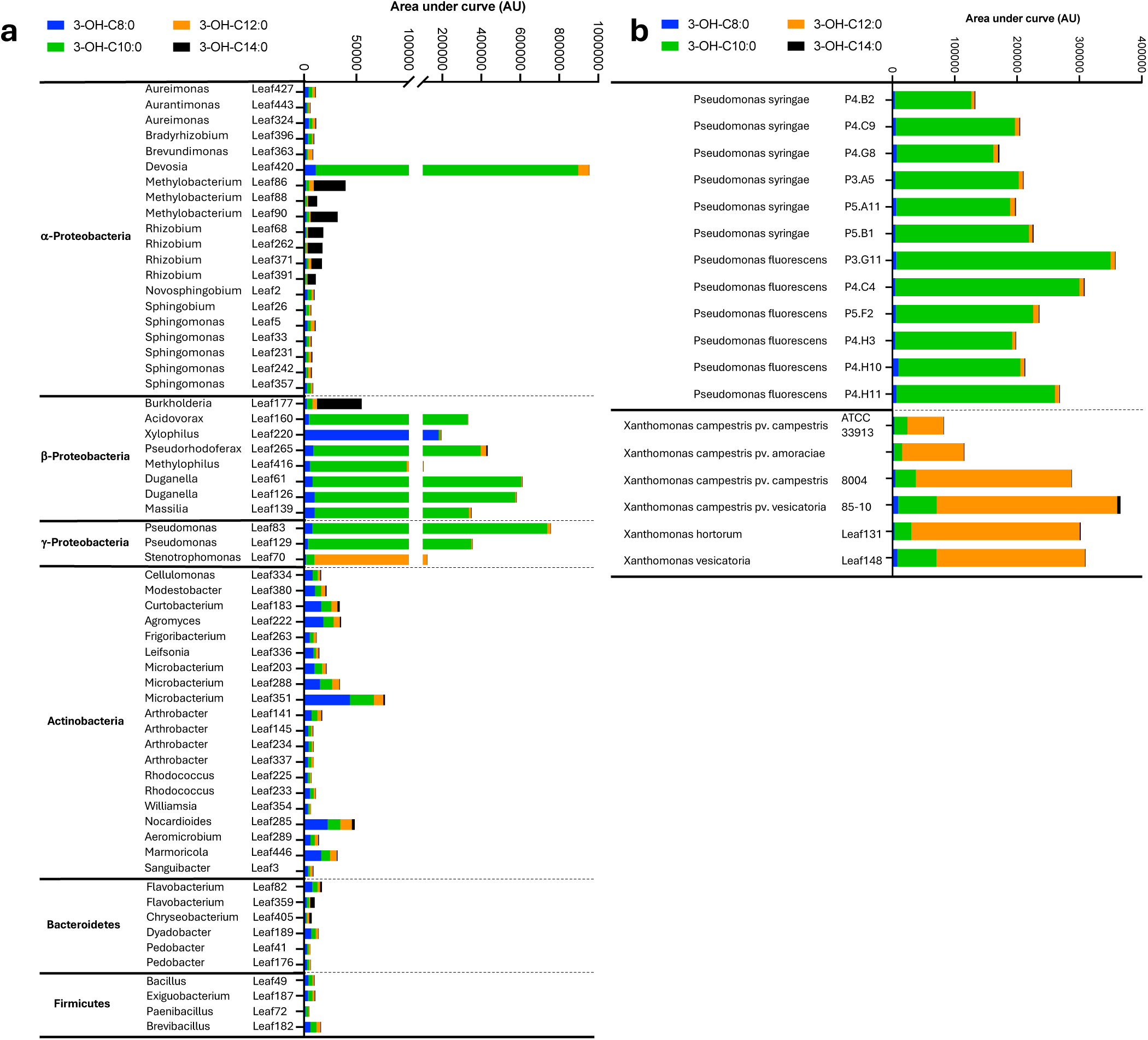
3-OH-FA content of different bacterial isolates from the natural *Arabidopsis thaliana* leaf microbiota and pathogenic Xanthomonas strains. **a, b** Content of 3-OH-FAs of the indicated chain length was quantified by LC-MS using internal standards, normalised to the bacterial dry weight, and is depicted as area under the curve. A representative result of one of two analyses is shown (see **Figure S10**).

## DISCUSSION

Previous studies established LORE as a PRR for bacterial mc-3-OH-FAs and related mc-HAAs and linked their LORE-dependent sensing to immunity in *A. thaliana* (Kutschera *et al*., 2019; Schellenberger *et al*., 2021; Eschrig *et al*., 2024b; Duchateau *et al*., 2025). Building on this, we show that LORE orthologs are conserved across all four major Brassicaceae lineages and that the LORE-like clade is absent from the sister family Cleomaceae. These findings place the origin of LORE after the Brassicaceae–Cleomaceae divergence and before the separation of the tribe Aethionemeae from core Brassicaceae, approximately 35–40 million years ago (Schranz & Mitchell-Olds, 2006; Hohmann *et al*., 2015). The functional complementation of *lore*-1 by *Aara*LORE from the basal lineage confirms that a functional mc-3-OH-FA receptor was already present at this early stage. LORE thus represents a Brassicaceae-specific innovation in innate immunity, the emergence of which coincides with a period of rapid ecological diversification and niche expansion within this family (Franzke *et al*., 2011; Hohmann *et al*., 2015; Huang *et al*., 2016).

Despite the broad conservation of *LORE* orthologs, several Brassicaceae species did not respond to mc-3-OH-FAs in our leaf-based ROS assay (**Fig. 1**). The 3-OH-FA-sensing function might have been lost in these species, as shown for *A. lyrata* and *A. halleri*, whose LORE orthologs lost the ability to homomerize (Eschrig *et al*., 2024b). Loss of PRR function within a plant family is a common phenomenon (Zhang *et al*., 2021; Fan *et al*., 2022; Snoeck *et al*., 2022). For LORE, the molecular mechanisms underlying loss of function can vary. In *A. lyrata* and *A. halleri*, LORE orthologs retain 3-OH-C10:0 binding but have lost the ability to homomerise, a prerequisite for signalling (Eschrig *et al*., 2024b). *Risl*LORE and *Bnig*LOREb likewise fail to complement *lore*-1 despite a functional, 3-OH-C10:0-binding ECD (**Fig. 2-4**), while the chimera are functional in *N. benthamiana*, suggesting defects located in the TMD-ICD region, possibly affecting homomerisation, intracellular signalling function or compatibility with the *A. thaliana* signalling network. Uncoupling of ligand binding from receptor activation represents a ‘silent receptor’ state that may reflect a stepwise process of receptor pseudo-functionalisation. However, because our species-level screening was limited to a single immune readout in leaf tissue, negative results should be interpreted with caution. Notably, for many species, accessions other than the sequenced ones were screened, and results cannot be directly correlated. In addition, species-specific differences in receptor expression patterns, tissue context, ligand accessibility in the apoplast or downstream signalling network activity may mask receptor-dependent responses. Indeed, several LORE orthologs from unresponsive species were functional upon heterologous expression, although the extent to which this activity reflects their functionality in the original species cannot be deduced from performance in the heterologous hosts. For RLP23, for instance, the origin of the C-terminal domain substantially influences compatibility with the SOBIR1 adapter kinase across species and thus signalling output (Yang *et al*., 2025). Nevertheless, the high proportion of LORE orthologs compatible with the *A. thaliana* signalling machinery suggests that, in addition to ligand binding, other signalling-relevant features are conserved across LORE orthologs from all Brassicaceae lineages.

Mc-3-OH-FA sensing has been conserved across the Brassicaceae family despite functional diversification of the chain-length preference. Hence, conservation of receptor-mediated sensing does not necessarily imply strict conservation of ligand selectivity. The chain-length preference patterns observed in our complementation and chimera experiments (**Fig. 3, 4**) were largely retained across heterologous expression systems, supporting the conclusion that chain-length preference is a receptor-intrinsic property encoded largely in the ECD. Elucidating the molecular basis of these differences through structural analysis or domain swaps among divergent orthologs will be an important next step.

All responsive LORE orthologs sense both 3-OH-C10:0 and 3-OH-C12:0, but they differ in the breadth of their chain-length preference profile (**Fig. 3, 4**). Some orthologs showed a narrow preference with strong 3-OH-C10:0 dominance, similar to *A. thaliana* Col-0, whereas others displayed a broader response profile with comparable sensitivity to 3-OH-C10:0 and 3-OH-C12:0, and in some cases also to 3-OH-C8:0 or 3-OH-C13:0. Several species, predominantly in lineage II, encode multiple LORE-like paralogs, but these do not appear to partition the ligand spectrum among them. This contrasts with the duplicated FLS2 paralogs in *Vitis riparia* and soybean, where one copy acquired expanded sensitivity to otherwise unrecognised flagellin variants from *Agrobacterium tumefaciens* (Fürst *et al*., 2020) or *Ralstonia solanacearum* (Wei *et al*., 2020), respectively. Likewise, the orthologous receptors SCORE in Citrus and CORE in Solanaceae sense closely related cold-shock proteins with distinct binding specificities (Ngou *et al*., 2025). In the LORE system, diversification occurred at the level of relative chain-length preference within a shared ligand class, with paralogs retaining overlapping specificity. Whether LORE paralogs differ in other properties, such as binding affinity, signalling activity, or tissue-specific expression patterns, remains to be determined.

Sensing of 3-OH-C10:0 is robust across all responsive species and LORE orthologs tested, indicating that this reflects the core function of this PRR. Sensing of other mc-3-OH-FAs therefore likely reflects species-specific fine-tuning of the receptor rather than a shift in primary specificity. Consequently, 3-OH-C10:0 is best suited as a ‘type elicitor’ of mc-3-OH-FAs for comparative studies across plant species and experimental systems. In contrast, no response to 3-OH-C14:0 was observed for any species or LORE-ortholog tested. 3-OH-C14:0 appears to be a cut-off length and may reflect biophysical constraints, because longer acyl chains reduce aqueous solubility and may be incompatible with receptor binding in the apoplastic environment, or may result from the absence of selective pressure to sense 3-OH-C14:0, given its low abundance in the bacteria profiled here (**Fig. 5**).

Our bacterial 3-OH-FA profiling reveals that the ligand range preferentially sensed by LORE orthologs centres on 3-OH-C10:0 and 3-OH-C12:0, the mc-3-OH-FAs most abundant in Proteobacteria, particularly β- and γ-Proteobacteria. Among these, *Pseudomonas* species are well known for producing diverse biomolecules comprising mc-3-OH-FAs, typically 3-OH-C10:0, including lipopolysaccharide, polyhydroxyalkanoates, rhamnolipids, HAAs, lipopeptides, and *N*-acyl-homoserine lactones (De Souza *et al*., 2003; Corsaro *et al*., 2004; Raaijmakers *et al*., 2006; Verlinden *et al*., 2007; Abdel-Mawgoud *et al*., 2010; Raaijmakers *et al*., 2010; Ranf *et al*., 2015; Gerster *et al*., 2022). *Pseudomonas* release free 3-OH-C10:0 during the synthesis of penta-acylated lipopolysaccharide in the outer membrane (Geurtsen *et al*., 2005; Ernst *et al*., 2006; Gerster *et al*., 2022). *Xanthomonas* and *Stenotrophomonas* produce lipopolysaccharide comprising 3-OH-C12:0 (Silipo *et al*., 2008; Casabuono *et al*., 2011; Naito *et al*., 2017). Both free mc-3-OH-FAs and mc-HAAs are part of the bacterial secretome and activate LORE-mediated immunity (Schellenberger *et al*., 2021; Duchateau *et al*., 2025). While bacteria do not necessarily produce free mc-3-OH-FAs as regular metabolites, these may be released from acyl carrier protein (ACP)- and coenzyme A (CoA)-bound mc-3-OH-FA intermediates, which are required in significant amounts for the synthesis of mc-3-OH-FA-containing biomolecules, through thioesterase activities (Zheng *et al*., 2004) or non-enzymatically upon cell lysis in the apoplast. Bacterial outer membrane vesicles may provide an additional delivery route for mc-3-OH-FAs (Katsir & Bahar, 2017). Since odd-numbered mc-3-OH-FAs are rare in nature, their biological relevance as elicitors remains to be determined; however, they are found in lipopolysaccharide, e.g. in *Xanthomonas* (Silipo *et al*., 2008; Casabuono *et al*., 2011), and polyhydroxyalkanoates (Ruth *et al*., 2007).

Notably, individual bacterial strains that produce high levels of mc-3-OH-FAs tend to accumulate predominantly one chain length rather than multiple chain lengths simultaneously at high abundance (**Fig. 5**). This suggests that the chain-length preference of a given LORE ortholog may determine which bacterial groups are most effectively detected: orthologs with narrow, 3-OH-C10:0-dominated preference profiles may be particularly suited to *Pseudomonas*-like producers, whereas orthologs with broader profiles may sense a wider range of Proteobacteria. The diversification of LORE chain-length preferences across Brassicaceae species could thus reflect adaptation to locally prevalent bacterial communities with distinct mc-3-OH-FA profiles. Moreover, mc-3-OH-FAs are not exclusive to bacteria. They are also produced by honeybees (Weaver *et al*., 1968) and leaf-cutter ants (Schildknecht & Koob, 1971). How broadly mc-3-OH-FAs are present in plant-interacting organisms, and whether non-bacterial sources are relevant to LORE-mediated immunity in Brassicaceae (Oldstone-Jackson *et al*., 2023), remains to be studied. The pathways by which different bacteria produce and release mc-3-OH-FAs during host interactions also require further investigation.

From an evolutionary perspective, the correspondence between LORE chain-length preferences and bacterial 3-OH-FA profiles is consistent with ecological matching rather than a classical co-evolutionary arms race. Unlike peptide MAMPs such as flagellin, whose epitopes can diversify under immune selection pressure (Cai *et al*., 2011; Mott *et al*., 2016; Colaianni *et al*., 2021; Parys *et al*., 2021), mc-3-OH-FAs are structurally constrained core metabolites: they are essential intermediates in bacterial lipid and lipopolysaccharide biosynthesis, and their chain-length distribution is determined primarily by core metabolic requirements rather than by selection to evade plant immune detection. This suggests that Brassicaceae LORE receptors have adapted to a relatively stable chemical landscape presented by their microbial environment, a scenario of asymmetric co-evolution in which the plant receptor diversifies while the microbial ligand remains under metabolic selection. The LORE/mc-3-OH-FA system thus provides an experimentally accessible case to study how ligand preference can diversify within a conserved PRR family when the ligand spectrum itself is biologically constrained.

To date, LORE has been shown to contribute to resistance against bacterial pathogens, as well as to systemic immunity triggered by beneficial root-associated bacteria and synthetic elicitor application (Kutschera *et al*., 2019; Schellenberger *et al*., 2021; Eschrig *et al*., 2024b; Duchateau *et al*., 2025; Li *et al*., 2025b). The natural LORE variants characterised here, with their distinct chain-length preference profiles and signalling outputs, provide a valuable resource for developing disease-resistant crops through breeding or PRR engineering (Boutrot & Zipfel, 2017; Ranf, 2018; Li *et al*., 2025a; Ngou *et al*., 2025; Yang *et al*., 2025). Since individual bacterial groups seem to typically produce predominantly one mc-3-OH-FA chain length (**Fig. 5**), selecting orthologs with a matching preference profile may enable tailored immune recognition of specific pathogenic or beneficial bacteria.

In conclusion, our results establish LORE as a Brassicaceae-specific PRR whose evolutionary trajectory can be traced from its origin to the present across all major lineages. Unlike PRR systems for which only presence–absence or on–off functionality has been analysed, the LORE system allows receptor evolution to be dissected at the levels of ligand chemistry, chain-length preference, and species-level immune responsiveness, within a biologically relevant and experimentally defined ligand landscape. This multi-level framework may serve as a model for analysing functional diversification in other PRR families in which receptor conservation is established, but the rules governing variation in ligand specificity remain unknown.

## MATERIALS AND METHODS

### Plant materials

*Arabidopsis thaliana* Col-0 was used as the primary genetic background in this study unless otherwise specified. *A. thaliana* Col-0^AEQ^, *lore*-1 knockout, the two complementation lines *p*At*LORE:*At*LORE*-CL1/*lore*-1 and *p*At*LORE:*At*LORE*-CL2/*lore*-1, and the p35S:At*LORE*/*lore*-1 overexpressing line (OE) all express cytosolic apoaequorin and were described previously (Ranf *et al*., 2015; Shu *et al*., 2021). The ecotypes of *A. thaliana*, other Brassicaceae, and non-Brassicaceae species used for ROS measurements are listed in **Tables S1 and S2**. Arabidopsis and other plants were grown in potting soil (ED73, Einheitserde, Germany) mixed with vermiculite (9:1) under short-day conditions (8 h light/16 h dark) at 22°C/20°C (day/night) (Hu *et al*., 2011; Briskine *et al*., 2017; Kiefer *et al*., 2019; Fernandez-Pozo *et al*., 2021). *N. benthamiana* and *Arabidopsis* plants used for seed propagation were grown under long-day conditions (16 h light/8 h dark) at 23°C/21°C (day/night).

### Bacterial strains

*Escherichia coli* DH5α was used for cloning, and *Agrobacterium tumefaciens* GV3101 (pMP90) for transformation of *A. thaliana* and *N. benthamiana*. Bacterial strains used for 3-OH-FA profiling are listed in **Table S3.**

### Elicitors for immune response assays

3-Hydroxydecanoic acid (3-OH-C10:0), 3-hydroxyoctanoic acid (3-OH-C8:0), 3-hydroxynonanoic acid (3-OH-C9:0), 3-hydroxyundecanoic acid (3-OH-C11:0), 3-hydroxydodecanoic acid (3-OH-C12:0), 3-hydroxytridecanoic acid (3-OH-C13:0), and 3-hydroxytetradecanoic acid (3-OH-C14:0) were purchased from Matreya LLC, USA. The proteinaceous elicitor flg22 (QRLSTGSRINSAKDDAAGLQIA) was synthesised by Pepmic, China.

### Screening of plants for 3-OH-FA-induced reactive oxygen species production

Leaf disks (4 mm in diameter) from 8- to 12-week-old *A. thaliana* and other plant species were floated on 200 μL of water in a 96-well white plate and incubated overnight in the dark. 30 minutes before measurement, the water was replaced with 100 μL solution containing 5 μM L-012 (FUJIFILM Wako Chemicals, Japan) and 2 μg/mL horseradish peroxidase (Roche, Switzerland). ROS levels were measured using a Luminoskan Ascent 2.1 microplate luminometer (Thermo Scientific, USA) for 10 minutes to obtain background signals, with readings taken at 1-minute intervals. After treatment with 25 μL of 5-fold concentrated elicitor solution, the ROS burst was monitored for 45 minutes. Water containing equivalent concentrations of MeOH or EtOH served as the negative control. Total luminescence was calculated by summing relative light units (RLU) from 1 to 45 minutes after elicitation. The maximum luminescence values between 1 and 45 minutes after elicitation were determined, normalised to the average luminescence during the 5 minutes prior to elicitation and MeOH control values obtained from separate leaf discs on the same plate were subtracted before plotting. Experiments were analysed using different luminometer plate readers, resulting in different ranges of relative light units (RLUs) depending on the reader.

### Gain-of-function assay for 3-OH-FA-induced reactive oxygen species production in Nicotiana benthamiana

LORE orthologs or chimera were expressed in leaves of 6-week-old *Nicotiana benthamiana* via *Agrobacterium*-mediated transformation. *A. tumefaciens* GV3101 (pMP90) carrying the expression constructs was grown on YEP agar supplemented with 50 μg/mL kanamycin, 30 μg/mL gentamicin and 100 μM acetosyringone at 28°C overnight. The bacterial culture was harvested and adjusted to an OD_600_ of 0.05 in infiltration buffer (10 mM MgSO₄, 10 mM MES, pH 5.5, 150 μM acetosyringone). The suspension was infiltrated into fully expanded *N. benthamiana* leaves using a needleless syringe. 3 days after infiltration, leaf disks (4 mm in diameter) were cut from the infiltrated leaf area and floated on 200 μL of water in a 96-well white plate overnight in the dark. 15-20 minutes before measurement, the water was replaced with 100 μL solution containing 200 μM luminol (Na salt; Roth, Germany) and 2 μg/mL horseradish peroxidase (Roth, Germany). ROS levels were measured using a Luminoskan Ascent 2.1 microplate luminometer (Thermo Scientific, USA) for 10 minutes to obtain background signals, with readings taken at 1-minute intervals. After treatment with 25 μL of 5-fold concentrated elicitor solution, the ROS burst was monitored for 60 minutes. Water containing equivalent concentrations of MeOH served as the negative control. For each leaf disk, luminescence values were normalised to the average luminescence during the 5 minutes prior to elicitation. Average MeOH control values, which were measured from separate leaf discs on the same plate, were subtracted, and total luminescence was calculated by summing relative light units (RLU) from 2 to 45 minutes after elicitation.

### Measurement of Arabidopsis cytosolic calcium ion elevations

*A. thaliana* cytosolic apoaequorin-expressing lines were cultured in half-strength MS liquid medium (2.2 g/L MS salts [Duchefa Biochemie, Netherlands], 2.5 g/L sucrose, 0.195 g/L 2-(N-morpholino)ethanesulfonic acid [MES], pH 5.7) under long-day conditions (16 h light/8 h dark) at 23°C/21°C (day/night) for 8–9 days. Seedlings were then transferred to a 96-well white plate and incubated overnight in 100 μL of water containing 5 μM coelenterazine-h (p.j.k. GmbH, Germany) in the dark. Luminescence signals were measured using a Luminoskan Ascent 2.1 (Thermo Scientific, USA) microplate luminometer. Background luminescence was recorded ten times at 10-second intervals prior to elicitation. Each seedling was treated with 25 μL of 5-fold concentrated elicitor solution, and luminescence was recorded every 10 seconds for 30 minutes after treatment. Water containing equivalent concentrations of MeOH or EtOH served as a negative control. To determine the total aequorin content of each seedling, 150 μL of discharge solution (2 M CaCl_2_, 20% EtOH) was added, and luminescence was recorded continuously for 2 min per well. Luminescence signals were normalised as L/L_max_ (luminescence counts per second/total luminescence counts) as described (Ranf *et al*., 2012), and total [Ca^2+^]_cyt_ signals were calculated by summing L/L_max_ values from 2 to 30 min after elicitation.

### Phylogenetic analysis of LORE orthologs and other SD-RLK

The sequences of LORE orthologs and other SD-RLKs were retrieved from public databases, and the protein sources, identities, and genome versions used for the phylogenetic analysis are listed in **Table S4.** Candidate protein sequences were analysed using SignalP-5.0 and TMHMM-2.0 to confirm the presence of signal peptides and TMDs, respectively (Krogh *et al*., 2001; Almagro Armenteros *et al*., 2019). Sequences lacking predicted signal peptides were further examined using RNA-seq data, and candidates with contigs covering missing N-terminal coding regions were manually corrected. For this study, ECDs were defined as extending from the signal peptide to approximately amino acids 410–440, encompassing conserved serine–glutamate motifs present in all LORE orthologs. The transmembrane and juxtamembrane regions were defined as extending from the serine–glutamate motifs to approximately amino acids 490–530, which include conserved lysine–leucine motifs. Intracellular domains were defined as extending from the lysine–leucine motifs to the glycine–arginine motifs adjacent to the stop codon. Protein sequences were aligned using the online tools MAFFT and MUSCLE available through the EMBL-EBI website. Phylogenetic trees were constructed using the MEGA XI software with the Neighbor-Joining method and a bootstrap analysis of 1,000 replicates. Evolutionary distances were calculated using the Poisson correction method (Tamura *et al*., 2021) embedded in MEGA XI. A phylogenetic tree without revealing bootstrap values was visualized using iTOL v7.

### Molecular cloning

The *A. tumefaciens* binary vector pGGPXhc was derived from pICSL4723 and redesigned with unique T-DNA regions for Golden Gate cloning. For mCherry fusion protein overexpression, the 35S promoter, LacZ cassette, mCherry, and heat shock protein terminator (tHSP) (Nagaya *et al*., 2010) were assembled within the T-DNA. pGGPXhcFR was generated by inserting a FastRed selection marker (Shimada *et al*., 2010) into the T-DNA. To express target genes under the At*LORE* promoter, the At*LORE* promoter, LacZ cassette, and tHSP were assembled in the T-DNA of pGGPXhcFR. To ensure compatibility with the pGGPXhc and pGGPXhcFR vectors, the CDS of LORE orthologs and other SD-RLKs were synthesised without Esp3I sites in coding sequences (Twist Biosciences). DNA fragments encoding ECDs, TMDs, and ICDs, with Esp3I sites at both ends, were synthesised separately. The native signal peptides of all ECDs were replaced with the *At*LORE signal peptide to optimise expression in *A. thaliana* and *N. benthamiana*, based on SignalP-5.0 predictions. Details of the synthetic DNA fragments are provided in **Table S5**. For ECD-mCherry fusion protein expression in *N. benthamiana*, ECD fragments were cloned into pGGPXhc-35S-LacZ-mCherry-tHSP by Esp3I-based Golden Gate cloning with blue–white screening. Full-length orthologs were obtained by assembling ECD, TMD, and ICD fragments of each ortholog, and chimera by assembling ortholog ECDs with AtLORE-TMD-ICD, which were then inserted into pGGPXhc-AtTCTP promoter-LacZ-terminator for expression in *N. benthamiana* or pGGPXhcFR-AtLORE promoter-LacZ-tHSP for *A. thaliana* transformation using the same cloning strategy.

### Expression and harvesting of LORE ortholog ECD proteins

Expression of recombinant LORE ECD proteins has been described previously (Shu *et al*., 2021; Shu *et al*., 2026). In brief, LORE ortholog ECD proteins were expressed in leaves of 6-week-old *Nicotiana benthamiana* via *Agrobacterium*-mediated transformation. *A. tumefaciens* GV3101 (pMP90) carrying the expression constructs was grown on LB agar supplemented with 50 μg/mL kanamycin and 100 μM acetosyringone at 28°C overnight. The bacterial culture was harvested, adjusted to an OD_600_ of 0.5 in infiltration buffer (10 mM MgSO₄, 10 mM MES, pH 5.5, 150 μM acetosyringone), and incubated at 28°C for 3 hours prior to infiltration. Cultures were mixed at equal volumes with *Agrobacterium* carrying the silencing suppressor p19 to a final OD₆₀₀ of 0.6. The suspension was infiltrated into fully expanded *N. benthamiana* leaves using a needleless syringe. Proteins secreted into the apoplast were harvested 3–4 days after infiltration. Detached leaves were submerged in Tris-buffered saline (TBS; 50 mM Tris base, 150 mM NaCl, pH 7.6) and vacuum-infiltrated to ∼26 in. Hg. After complete infiltration, excess surface buffer was removed, and leaves were rolled and placed into a 30 mL syringe barrel positioned inside a 50 mL tube. Samples were centrifuged at 800 × g for 20 minutes at 4°C to collect apoplastic washing fluid (AWF), which was further cleared by centrifugation at 8,000 × g for 20 minutes at 4°C. Prior to ligand depletion assays, AWF was desalted using PD-10 columns (Cytiva, USA) and concentrated to the desired protein concentration using Vivaspin 20 centrifugal filters (30,000 MWCO; Sartorius, Germany). Expression of LORE ECD–mCherry fusion proteins was confirmed by SDS–PAGE and western blotting using an anti-RFP antibody (ChromoTek, Germany).

### Ligand depletion assay

The ligand depletion assay was described previously (Shu *et al*., 2021; Shu *et al*., 2026). In brief, proteins in concentrated *N. benthamiana* AWFs were adjusted to the indicated concentrations with water. Protein samples were mixed with elicitors at a 9:1 (v:v) ratio, with elicitors prepared in water containing less than 1% MeOH, and incubated for 1 hour at 4°C on a rotator. Unbound ligands were separated from the mixture using Vivaspin 500 centrifugal filters with a 30,000 MWCO (Sartorius, Germany) and were collected in the filtrate. The ligands in the filtrates were detected by [Ca^2+^]_cyt_ measurements using LORE-OE and *lore*-1 seedlings.

### Generation of transgenic Arabidopsis

Transgenic Arabidopsis plants were generated using the floral dip method (Logemann *et al*., 2006). *A. tumefaciens* GV3101 was transformed with pGGPXhcFR binary vectors containing different LORE orthologs under control of the *AtLORE* promoter (Ranf *et al*., 2015) and the FastRed selection marker (Shimada *et al*., 2010) (**Table S6**). *A. tumefaciens* was grown on LB agar for two days, then washed and suspended in the solution containing 5% sucrose and 0.02% Silwet-77 to an OD_600_ of 2.0. *lore*-1 flowers were dipped in the suspension for 30-60 seconds and maintained in high humidity overnight. The transformed TagRFP-expressing seeds were screened using a stereomicroscope equipped with Fluorescent Adaptor, Green set (Excitation: 510-540 nm, emission filter: 600 nm longpass) (Nightsea, USA).

### RT-PCR

Total RNA from 9-day-old Arabidopsis seedlings grown in half-strength MS liquid medium was extracted according to the procedure of the Direct-zol RNA Mini Prep kit (Zymo Research) with TRIzol® Reagent (Thermo Fisher Scientific, USA). cDNA was synthesized by RevertAid RT Reverse Transcription Kit (Thermo Fisher Scientific, USA), and the manufacturer’s instruction was followed. Semi-quantitative PCR was performed with SupraTherm Taq Polymerase (Genecraft, Germany) under the following conditions and primers (**Table S7**): 95°C for 1 minute, then 28 cycles (*EF1-alpha*) or 36 cycles (*LORE*-ortholog) of 95°C for 10 seconds, 54°C for 10 seconds, and 72°C for 45 seconds, followed by 72°C for 5 minutes.

### 3-OH-FA profiling in bacteria

*At-*LSPHERE and other bacterial isolates (**Table S3**) were grown on R2A agar medium supplemented with 0.5% (v/v) MeOH at 28°C for five days. Bacteria were collected from the plates and dissolved in ice-cold ddH_2_O. Cells were harvested by centrifugation (3,000 × g, 10 min, 4°C) and washed twice with ice-cold ddH_2_O. Bacterial pellets were lyophilised. Crude fatty acid extraction was performed following the lipid extraction method of Bligh and Dyer (1959), with modifications. In brief, 50 mg of freeze-dried cells were incubated four times overnight in 3 mL MeOH/CHCl_3_ with alternating concentrations (2:1 (v/v) and 1:2 (v/v)) at 4°C. Solvents were evaporated by using a rotary evaporator at 40 mbar and a water bath (40°C). Finally, extracts were dissolved in 3 mL 1:2 (v/v) MeOH/CHCl_3_ and added to 0.5 mL ddH_2_O. After centrifugation (3,000 × g, 10 min, 4°C), the non-polar and polar phases were collected separately and freeze-dried. For the liquid chromatography-mass spectrometry (LC-MS) analysis, 50 µL of this extraction mixture was resolved in 50 µL of MeOH. UHPLC-MS/MS analysis was performed as described by (Kutschera *et al*., 2019). Absolute amounts of 3-OH-C8:0, 3-OH-C10:0, 3-OH-C12:0, and 3-OH-C14:0 are depicted as the area under the curve, arbitrary units (AU). For quantitative comparison, the area under the curve values from LC-MS analysis were normalised to the corresponding extract weights obtained from 50 mg of lyophilised cells.

## ACKNOWLEDGEMENTS

We thank Klaas Bouwmeester (Wageningen University, Netherlands), Uwe Conrath (RWTH Aachen, Germany), Pascal Falter-Braun (Technical University of Munich, Germany), Ralph Hückelhoven (Technical University of Munich, Germany), Thomas Mitchell-Olds (Duke University, USA), Marcel Quint (Martin Luther University Halle-Wittenberg, Germany), Stefan A. Rensing (University of Freiburg, Germany), Ulrich Schaffrath (RWTH Aachen, Germany), H.A. van den Burg (University of Amsterdam, Netherlands), Julia Vorholt (ETH Zurich, Switzerland), and Detlef Weigel (Max Planck Institute for Biology, Tübingen, Germany) for providing plant and bacterial material. This study received funding from the German Research Foundation: Priority Program SPP 2125 DECRyPT (Deconstruction and Reconstruction of the Plant Microbiota), subproject No. 401867691 to C.D. and No. 401843486 to S.R., and Collaborative Research Centre CRC924, subproject No. B10 to S.R., and the Swiss National Science Foundation: Grant No. 310030_208139/1 to S.R.

---

**Figure S1.**
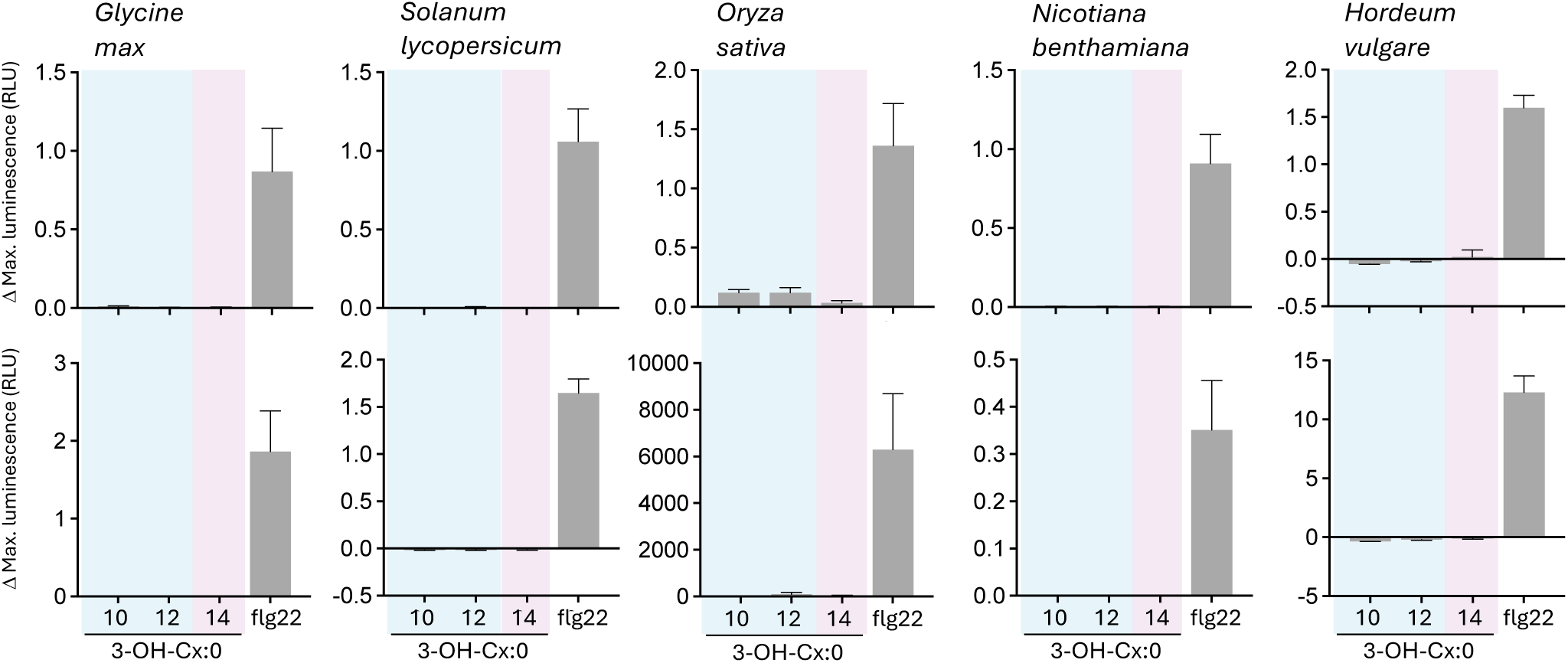
Response to 3-OH-FAs of different chain length in different plant species. Maximal ROS accumulation above MeOH control level in leaf disks of the indicated species upon elicitation with 5 µM 3-OH-FAs of the indicated chain length or 500 nM flg22 (mean ± SD, n = 8). Results from two independent experiments per species are shown in the upper and lower panel.

**Figure S2.**
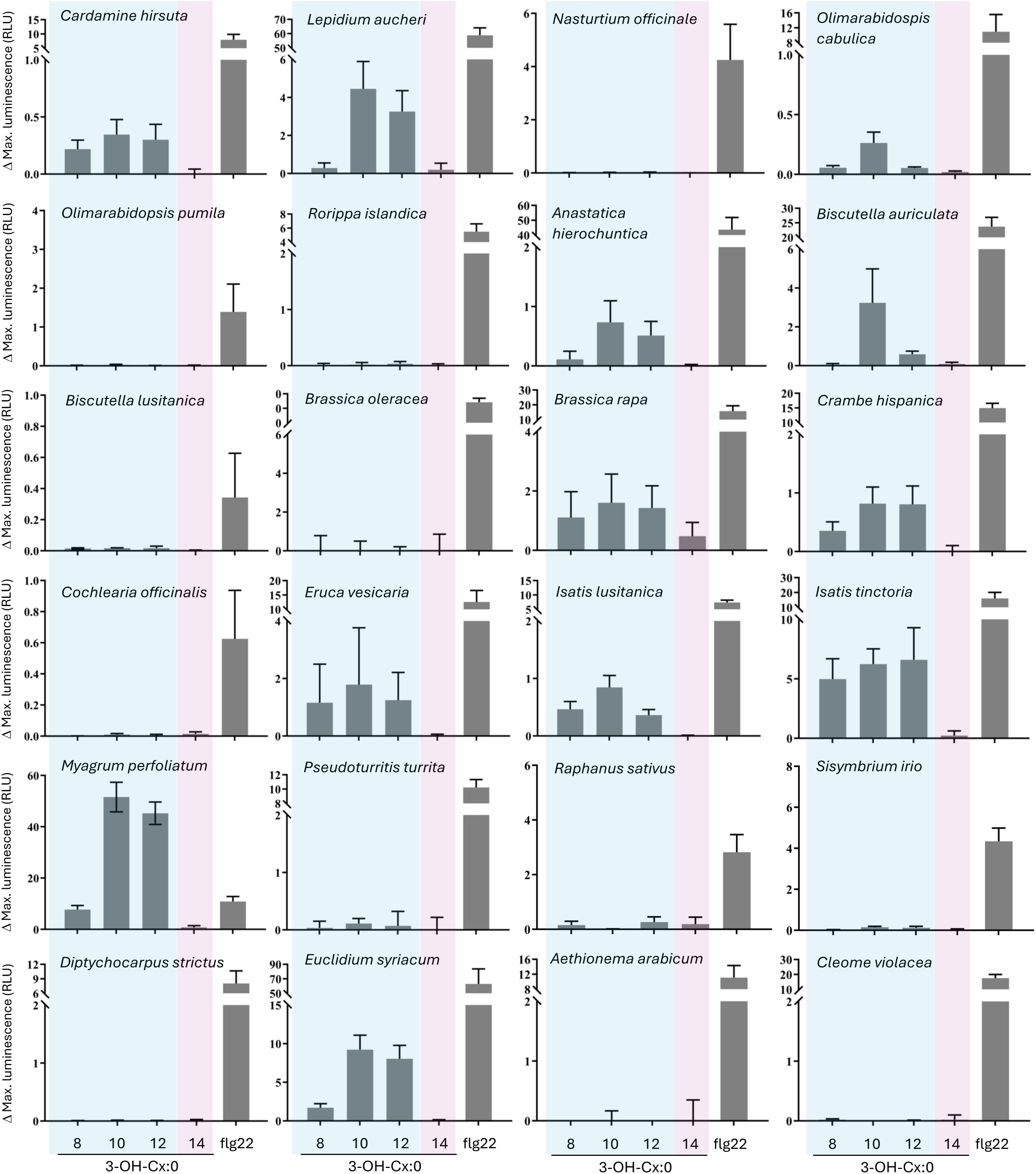
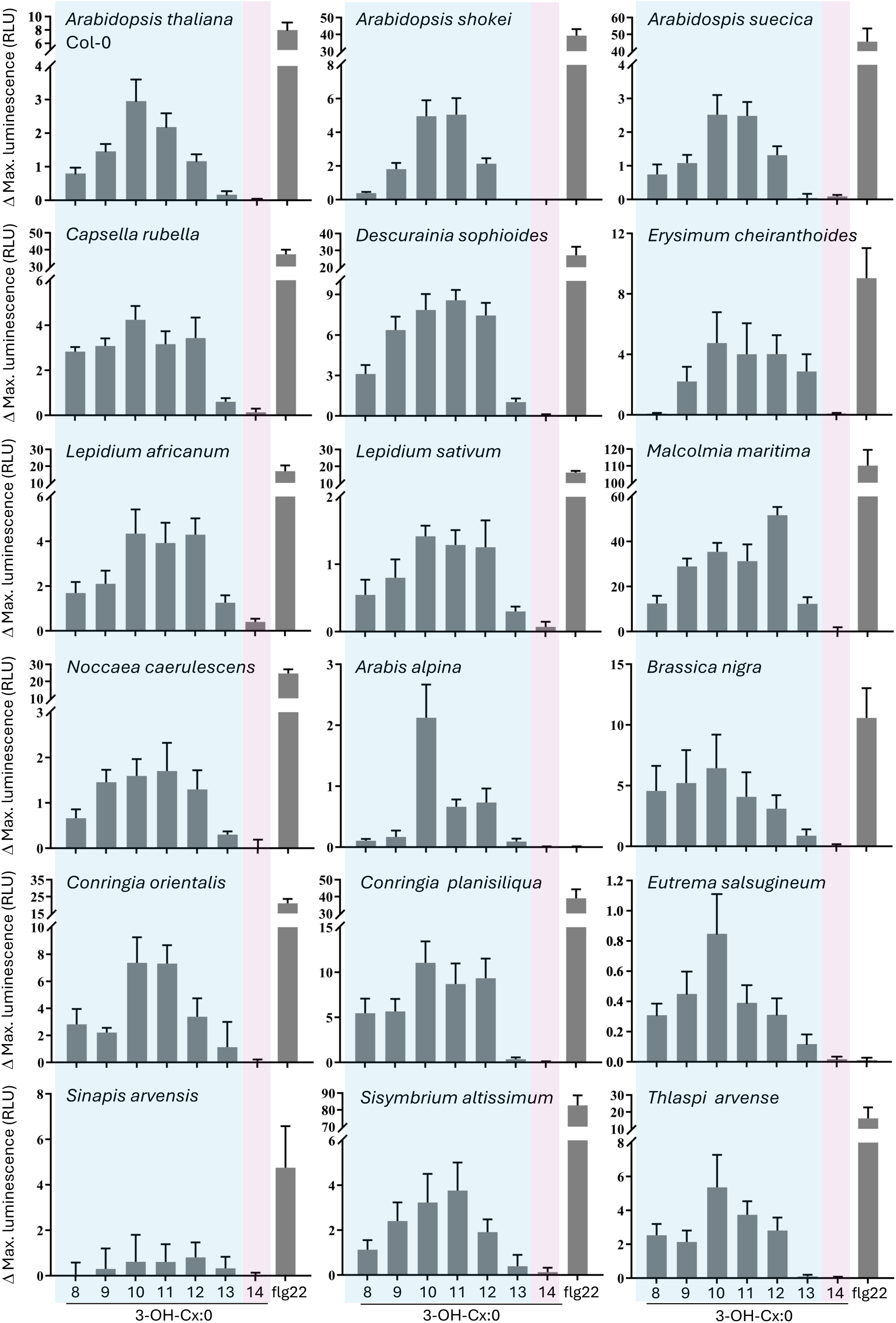
Diversity of 3-OH-FA chain-length sensing in Brassicaceae species and Cleomaceae outgroup. Maximal ROS accumulation above MeOH control level in leaf disks of the indicated Brassicaceae species upon elicitation with 5 µM 3-OH-FAs of the indicated chain length or 500 nM flg22 (mean ± SD, n = 8). Two or three independent experiments yielded similar results.

**Figure S3.**
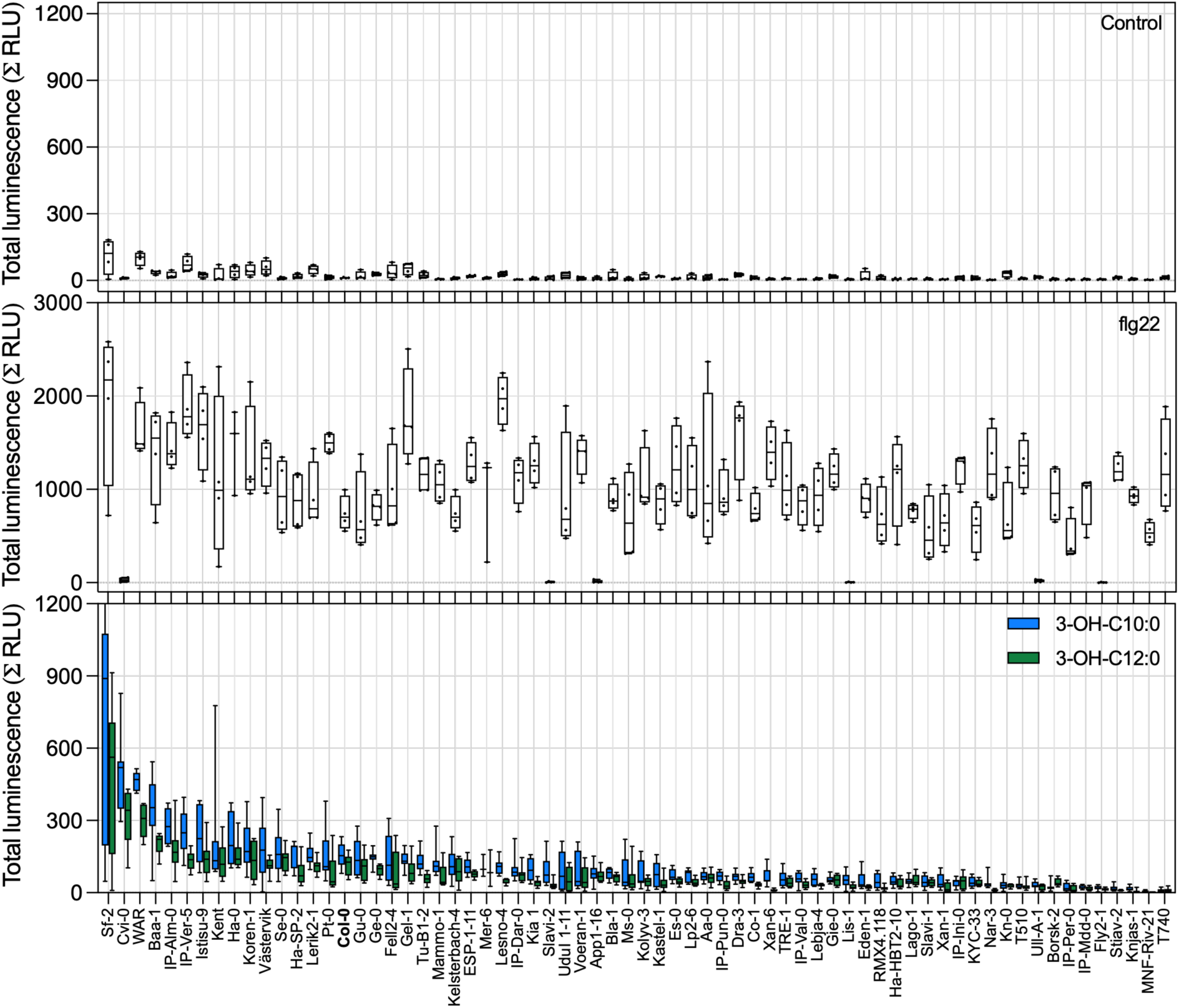
Sensitivity to 3-OH-C10:0, 3-OH-C12:0, and flg22 in leaves of Arabidopsis accessions. 68 accessions were tested for production of ROS in leaf disks from 9-week-old plants upon treatment with 1 µM 3-OH-C10:0 (n = 8), 1 µM 3-OH-C12:0 (n = 8), 100 nM flg22 (n = 4), or MeOH as a control (n = 4). Luminescence (relative light units, RLU) was measured at 1-minute intervals for 45 minutes after treatment and summed. Total luminescence in each sample is plotted as box and whiskers plots (Center lines are medians; boxes extend from 1^st^ quartile to 3^rd^ quartile; whiskers plot up to maximal values and down to minimal values).

**Figure S4.**
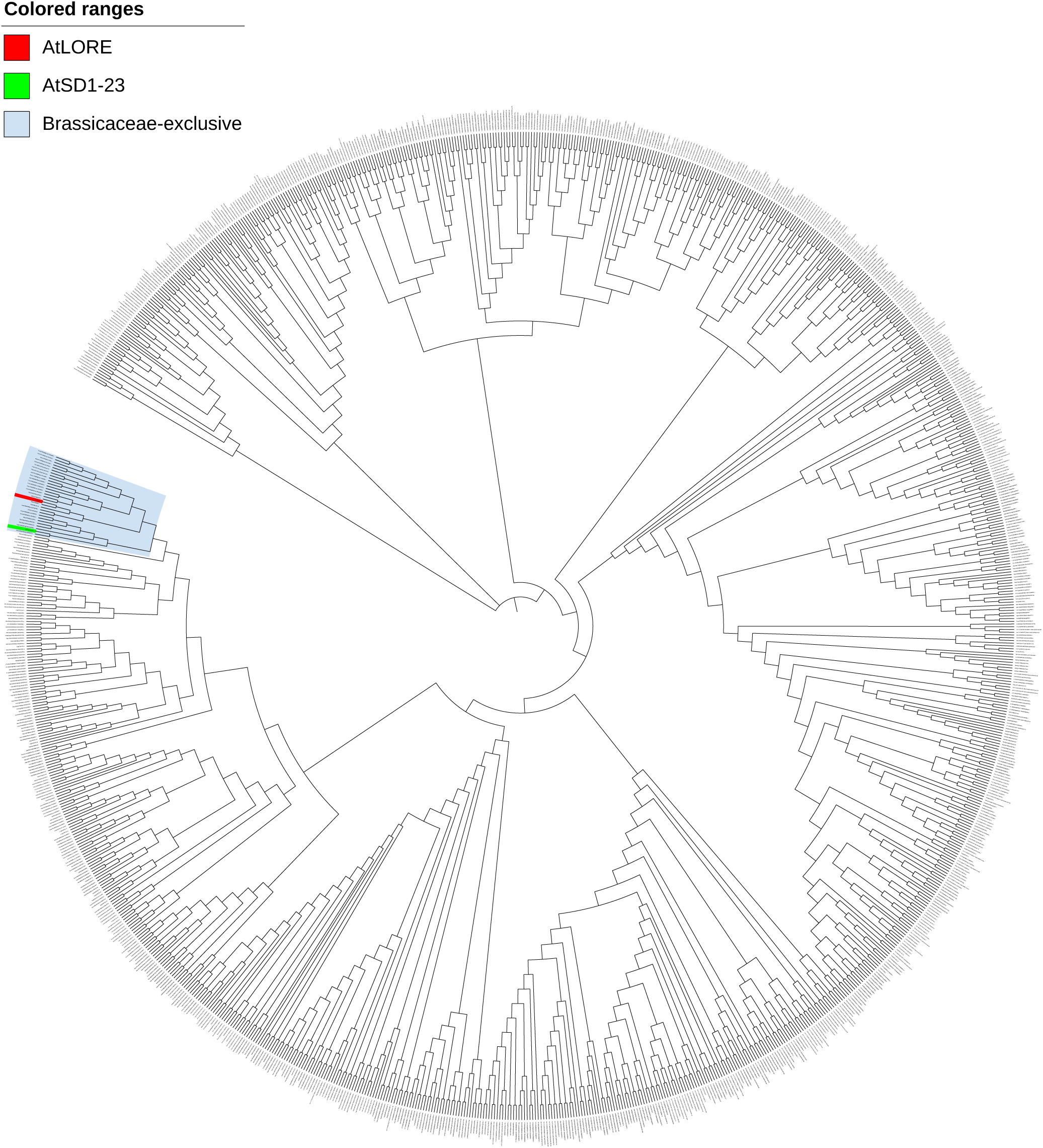
Phylogenetic analysis of LORE-related SD-RLKs in plants. The top 1000 closely related LORE-like genes, retrieved from genome sequences of 20 species from the Phytozome database, were aligned using MUSCLE. Phylogeny was analyzed in MEGA XI using the Neighbor-Joining method and visualized with iTOL v7. These 20 species include *Arabidopsis thaliana, Capsella rubella, Myagrum perfoliatum, Brassica oleracea, Eruca vesicaria, Thlaspi arvense, Brassica rapa, Eutrema salsugineum, Cleome violacea, Carica papaya, Theobroma cacao, Phaseolus vulgaris, Fragaria vesca, Vitis vinifera, Solanum tuberosum, Solanum lycopersicum, Coffea arabica, Beta vulgaris, Sorghum bicolor, Zea mays, Brachypodium distachyon, Oryza sativa, Musa acuminata,* and *Citrus clementina*. *At*LORE (red) and *At*SD1-23 (green) are in a Brassicaceae-exclusive lineage (blue).

**Figure S5.**
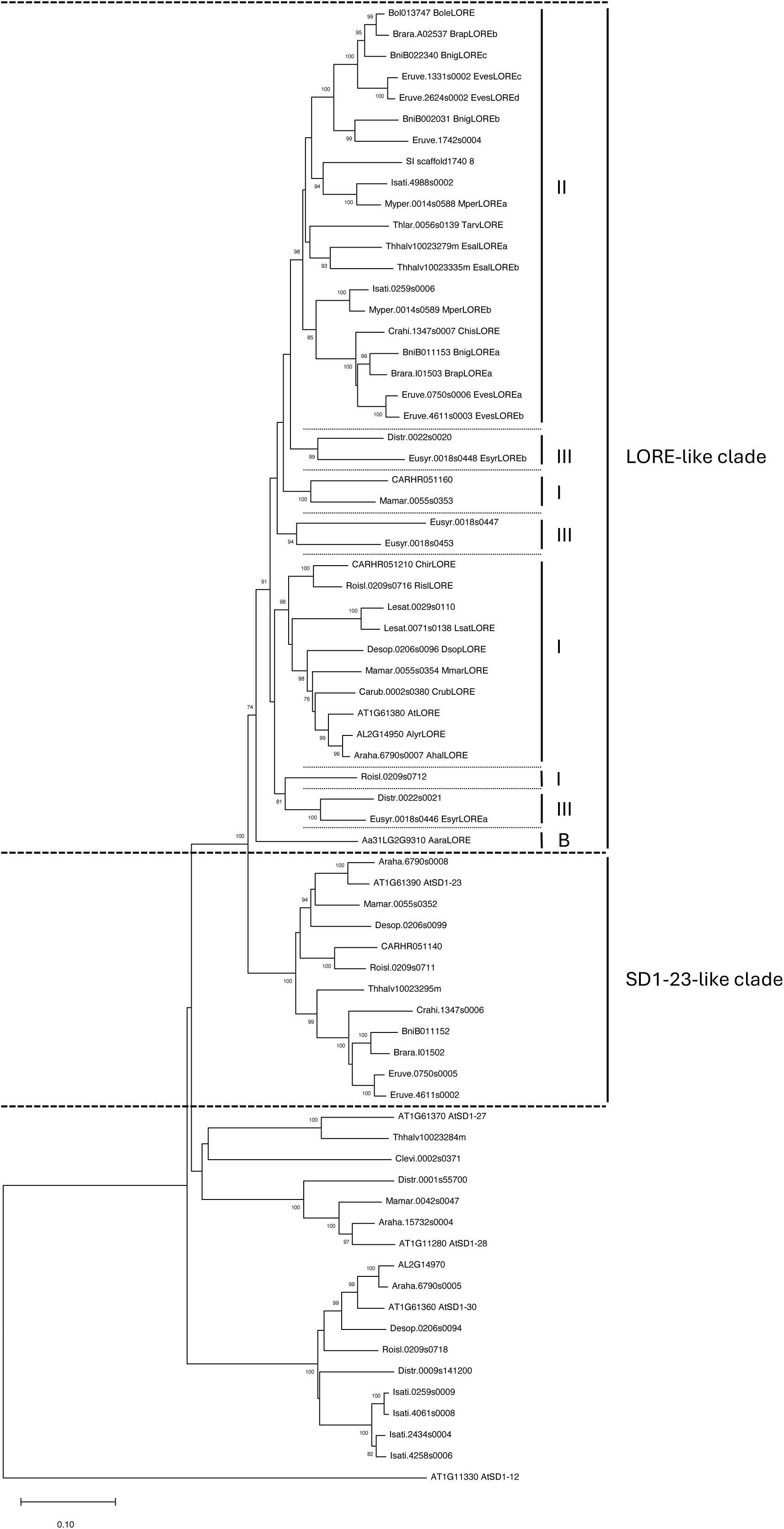
Phylogeny of LORE-like receptors and related SD-RLKs. 70 candidate protein sequences (**Table S2**) were aligned using the MUSCLE algorithm. Phylogeny was analysed in MEGA XI using the Neighbour-Joining method with bootstrap support (1000 replicates). Bootstrap values over 70 are shown next to branches. Scale bar indicates the evolutionary distance computed using the Poisson correction method. I, II, III and B (basal) indicate the lineages of the Brassicaceae species.

**Figure S6.**
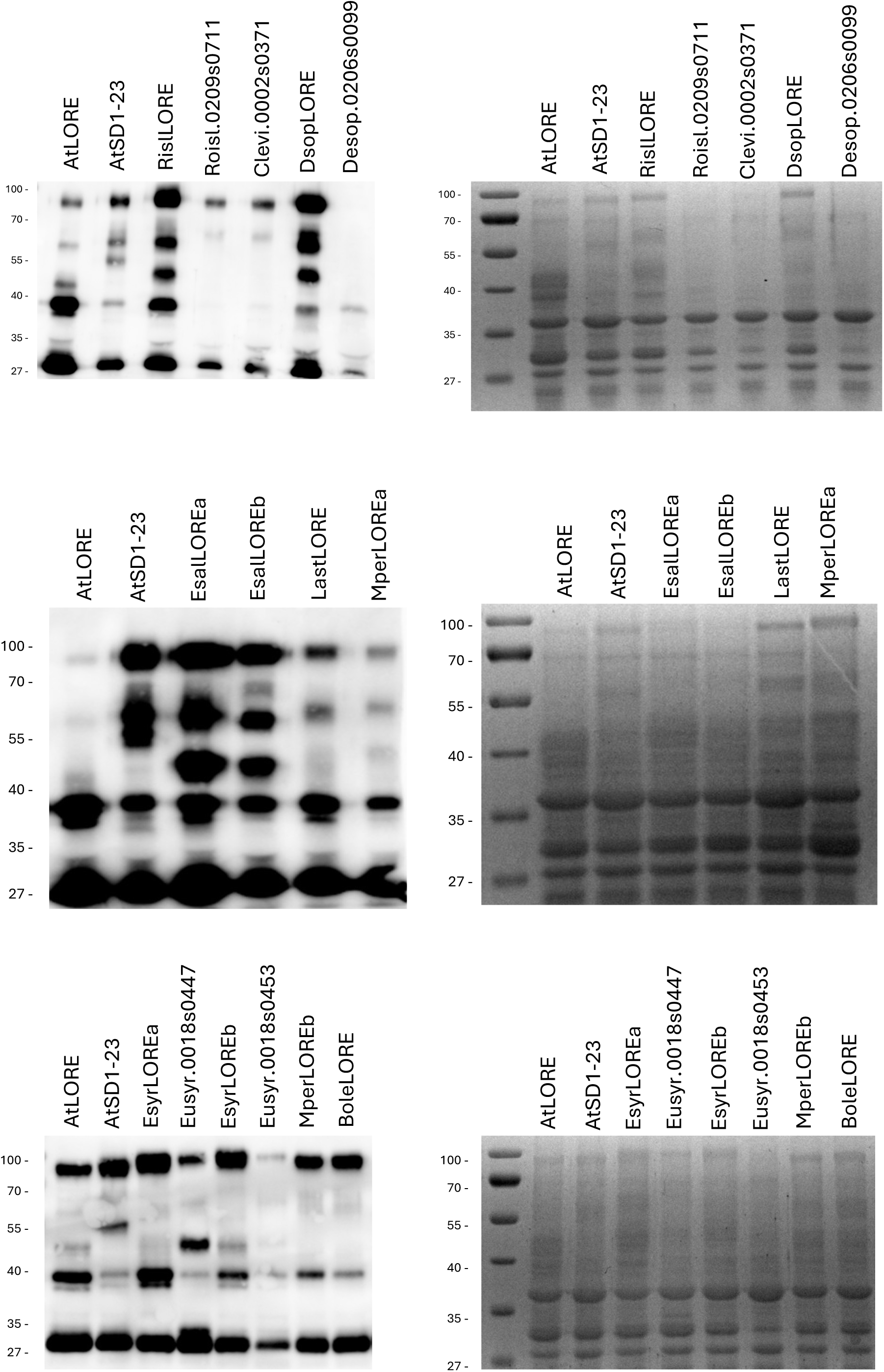
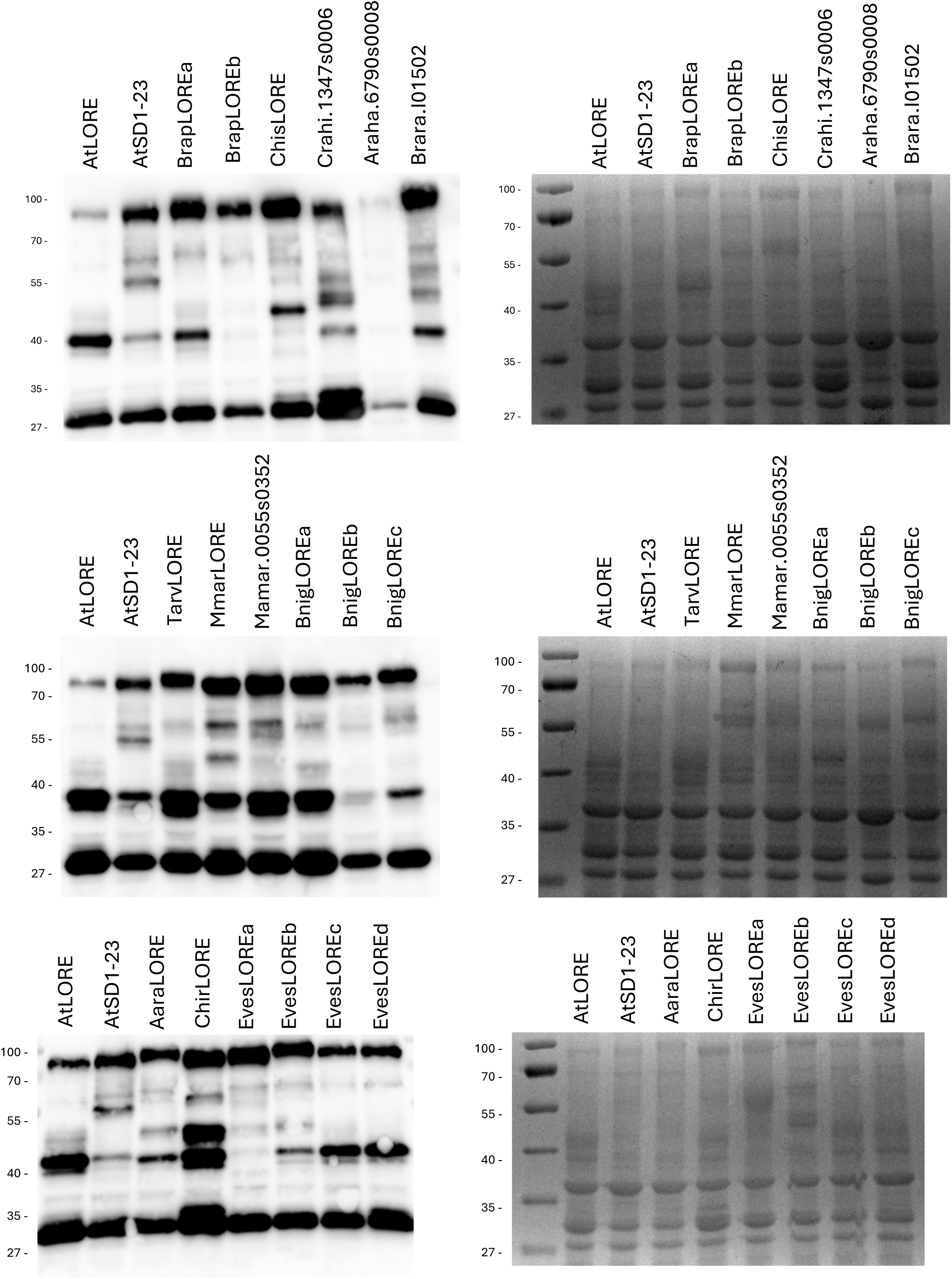
Anti-RFP immunoblot of desalted and concentrated AWFs from *N. benthamiana* leaves expressing different SD-RLK ECD-mCherry fusion proteins. Left panel: anti-RFP immunoblot, right panel: Coomassie Brilliant Blue (CBB) staining of SDS-polyacrylamide gel. AWFs were harvested 5 days after Agrobacterium infiltration in *N. benthamiana.* Desalted and concentrated AWFs corresponding to 7.5 !g total protein were loaded. The calculated molecular weights of SD-RLK ECD-mCherry fusions are 70-80 kDa. The loading of the samples was confirmed by Coomassie Brilliant Blue (CBB) staining of the SDS-polyacrylamide gel prior to blotting. One representative immunoblot of two experiments is shown.

**Figure S7.**
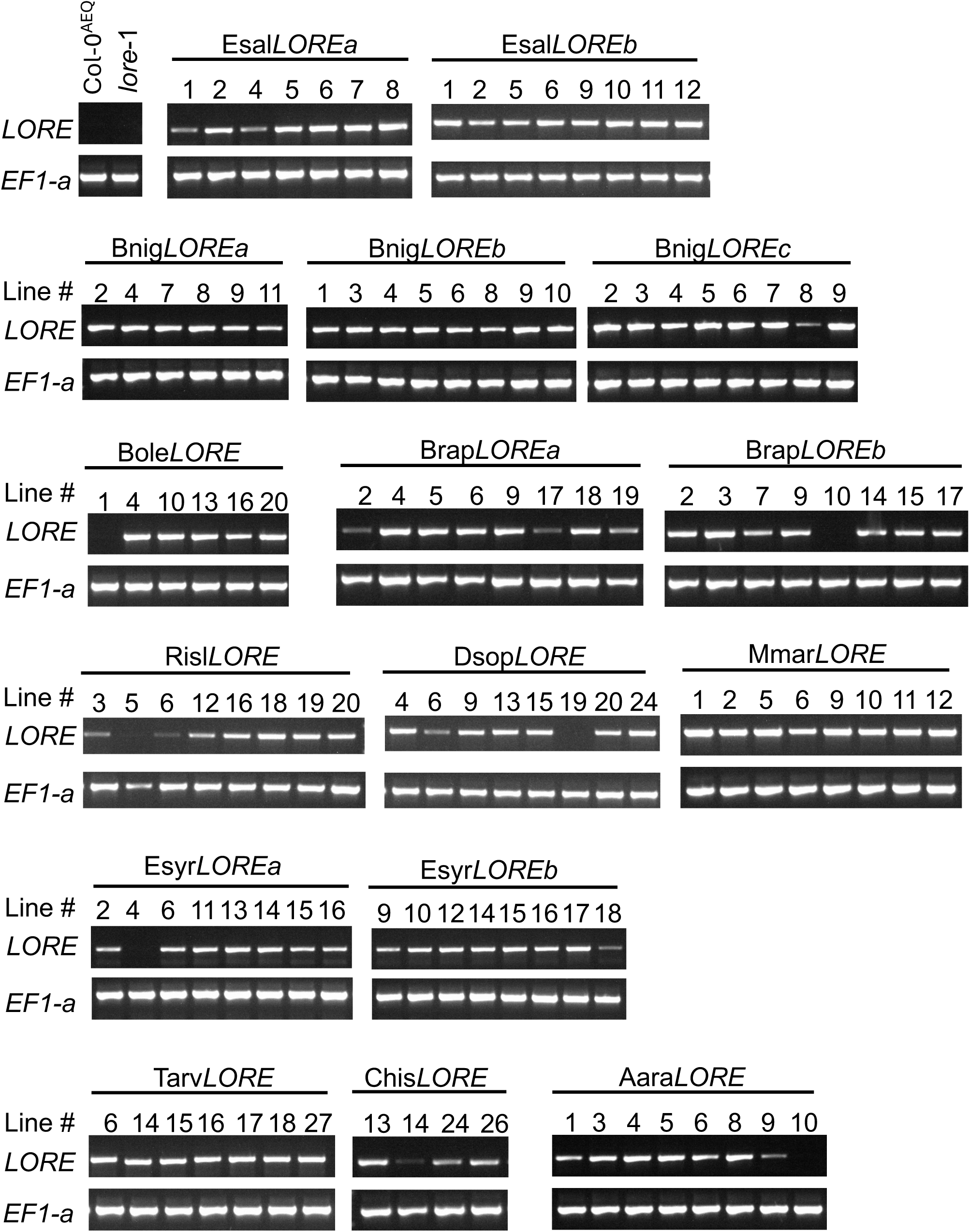
Expression of *LORE* orthologs in complementation lines. The expression of transgenes in *lore-*1 was analysed by RT-PCR with transgene-specific primers and analysed by agarose gel electrophoresis. Col-0^AEQ^ and *lore-*1 served as negative controls. The numbers above each lane indicate independent transgenic lines. *EF-1*α was used as an internal control of RT-PCR. Primer sequences are listed in **Table S7**.

**Figure S8.**
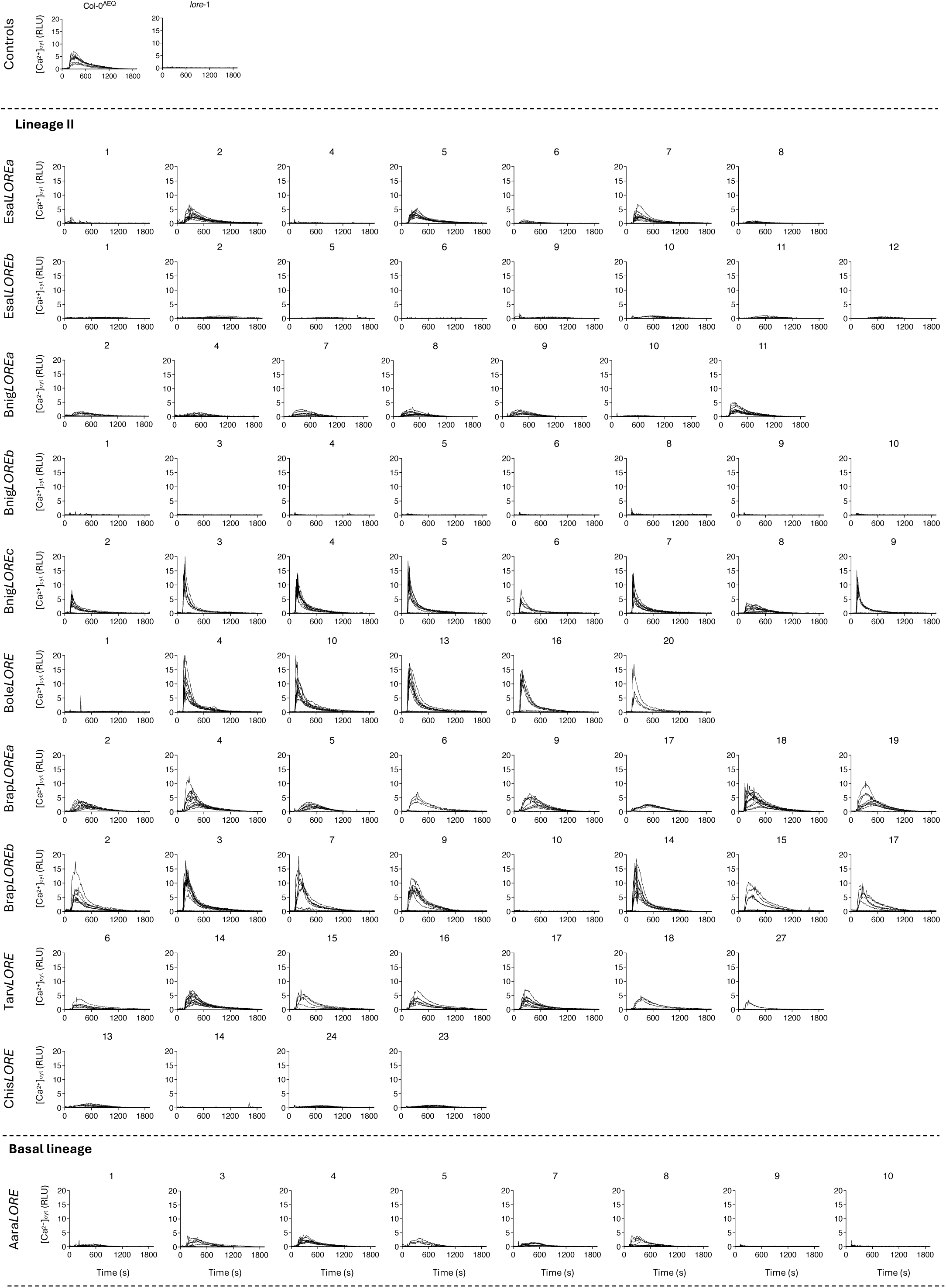
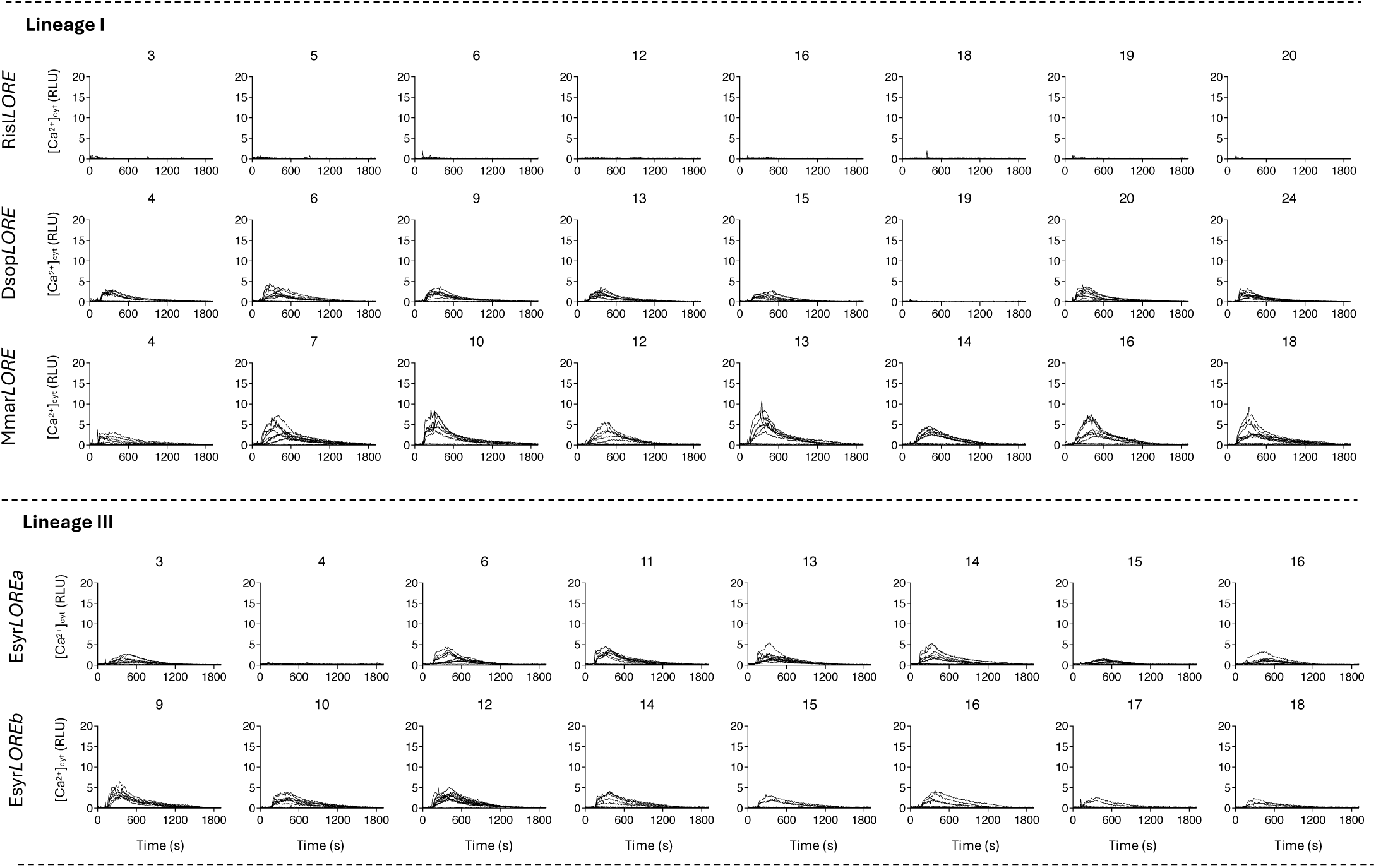
3-OH-FA-triggered immune responses in *Arabidopsis thaliana lore*-1 lines expressing various *LORE* orthologs spanning all four Brassicaceae lineages. [Ca^2+^]_cyt_ kinetics were measured in *LORE* ortholog-complemented *A. thaliana lore*-1 seedlings after elicitation with 1 µM 3-OH-C10:0. The names of *LORE* orthologs indicate the expressed transgenes, and the number above each kinetics graph indicates an independent transgenic line. Col-0^AEQ^ and *lore*-1 seedlings were included as positive and negative controls, respectively. Segregating T2 populations of independent lines were analysed, with 12 individual seedlings randomly selected from each population. The kinetics of individual seedlings from a single T2 population are shown in the same graph, and one of two independent experiments with the same pattern is shown. Proof of transgene expression for all complementation lines is provided in **Figure S7**.

**Figure S9.**
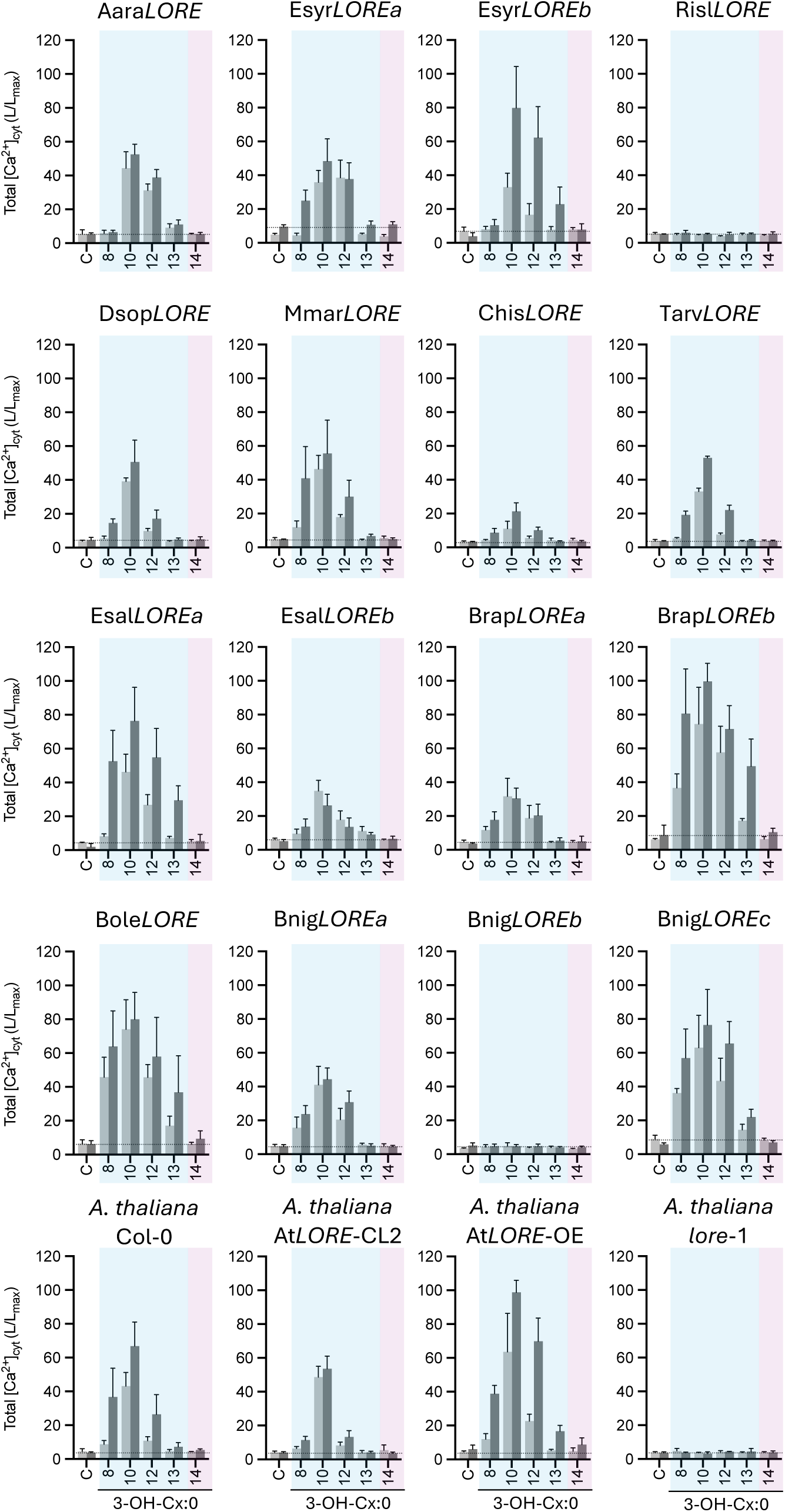
3-OH-FA chain-length preference of LORE-like receptors expressed in *Arabidopsis thaliana lore-*1. Total [Ca^2+^]_cyt_ responses were measured in *LORE* ortholog-complemented *A. thaliana lore*-1 seedlings after elicitation with 1 µM (light grey) or 5 µM (dark grey) 3-OH-FAs of the indicated chain length or MeOH as control (C). Dashed lines indicate average total [Ca^2+^]_cyt_ levels of controls as reference; shaded areas indicate medium (blue) or long (magenta) chain-length range of 3-OH-FAs. The T3 generations analysed were homozygous (based on FastRed), or, for heterozygous lines, FastRed-positive seeds were selected. One of two independent experiments with the same pattern is shown for 5 µM and one experiment for 1 µM (mean ± SD; n ≥ 4 seedlings per ortholog and chain length). Replication of Figure 3 from a set of independent transgenic lines. CL, complementation line; OE, overexpressing line.

**Figure S10.**
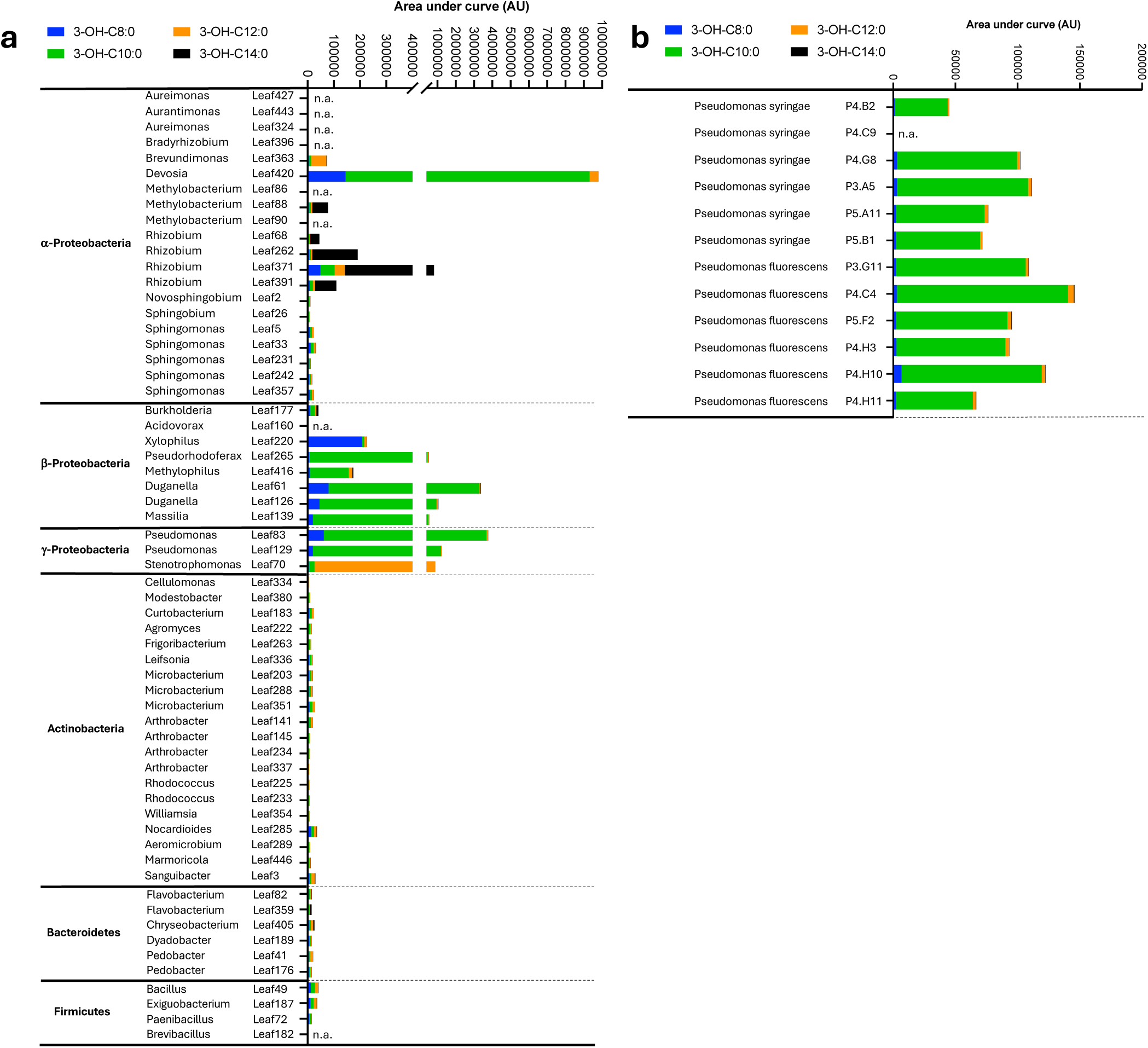
3-OH-FA content of different bacterial isolates from the natural *Arabidopsis thaliana* leaf microbiota. **a, b** Content of 3-OH-FAs of the indicated chain length was quantified by LC-MS using internal standards, normalised to the bacterial dry weight, and is depicted as area under the curve. Independent replication of the analysis shown in Figure 5. n.a., not analysed.

